# A Multi-Organ Single-Cell Atlas Maps Allergen and Organ-Specific Cellular Dynamics in the Progression from Food Allergy to Anaphylactic Shock

**DOI:** 10.64898/2026.02.04.703914

**Authors:** Richard L. Jayaraj, Xue Xia, Mahmoud Al-Azab, Yiyiao Liang, Sohaib H. Mazhar, Seth L. Alpert, Saima Khan, Amina Nazir, Jiawen Xu, Chengli Li, Bing Huang, Mourad Aribi, Xin Li, Abdelouahab Bellou

**Affiliations:** Institute of Sciences in Emergency Medicine, Department of Emergency Medicine, Guangdong Provincial Peoples Hospital, Guangdong Academy of Medical Sciences, Guangzhou 510080, China; Guangdong Cardiovascular Institute, Guangdong Provincial People’s Hospital, Guangdong Academy of Medical Sciences, Guangzhou 510080, China; Henan Key Laboratory for *Helicobacter pylori* and Digestive Tract Microecology, The Fifth Affiliated Hospital of Zhengzhou University and State Key Laboratory of Metabolic Dysregulation & Prevention and Treatment of Esophageal Cancer, Zhengzhou 450000, China; Department of Medical Microbiology, Faculty of Medicine, University of Science and Technology, Aden, Yemen; School of Basic Medical Sciences, Guangzhou Medical University, Guangzhou; Division of Nephrology and Vascular Biology Research Center, Beth Israel Deaconess Medical Center and Department of Medicine, Harvard Medical School, Boston, MA 02115, USA; Guangdong Provincial Key Laboratory of Gastroenterology, Department of Gastroenterology, Nanfang Hospital, Southern Medical University, Guangzhou, Guangdong, China; Laboratory of Applied Molecular Biology and Immunology, Abou-Bekr Belkaid University of Tlemcen, Tlemcen, 13000, Algeria; China-Algeria Joint Laboratory on Emergency Medicine and Immunology, Guangzhou 510080, China; Department of Emergency Medicine, Guangdong Provincial People’s Hospital (Guangdong Academy of Medical Sciences), Guangzhou 510080, China; Department of Emergency Medicine, Wayne State University, Detroit, MI, USA; Global Network on Emergency Medicine, Brookline, MA, USA

## Abstract

The transition from food allergic sensitization to life-threatening anaphylactic shock (AS) with complex multi-organ cellular interactions has been poorly understood. We herein conducted a multi-organ single-cell sequencing of heart, lung, and intestinal tissues from mouse models of food allergy, sensitized to bovine serum albumin (BSA) or ovalbumin (OVA), across both sensitization (Phase I) and anaphylactic shock (Phase II). We found that organ-specific pathologies included cardiac vascular dysfunction with pericyte loss and cardiomyocyte adaptation, distinct lung responses (Th2/B-cell predominance with BSA vs. neutrophilia with OVA), and profound intestinal remodeling featuring Paneth cell expansion. We also identified conserved inflammatory pathways (e.g., NOD-like receptor, NF-κB, JAK-STAT signaling) and unique organ specific signatures. Notably, phase-dependent reprogramming showed NLRP3 inflammasome activation and suppressed anti-inflammatory responses during AS. Cell-cell communication networks shifted from stromal-immune crosstalk in Phase I to dominant myeloid-vascular/neuroimmune interactions in Phase II. We further linked early extracellular matrix remodeling to late-stage neuromodulatory responses in Schwann and smooth muscle cells. This multi-organ atlas elucidates the coordinated, organ-selective immune cascade driving FA to AS progression, providing a foundational resource for developing targeted therapeutic strategies.

**Graphical Abstract:** 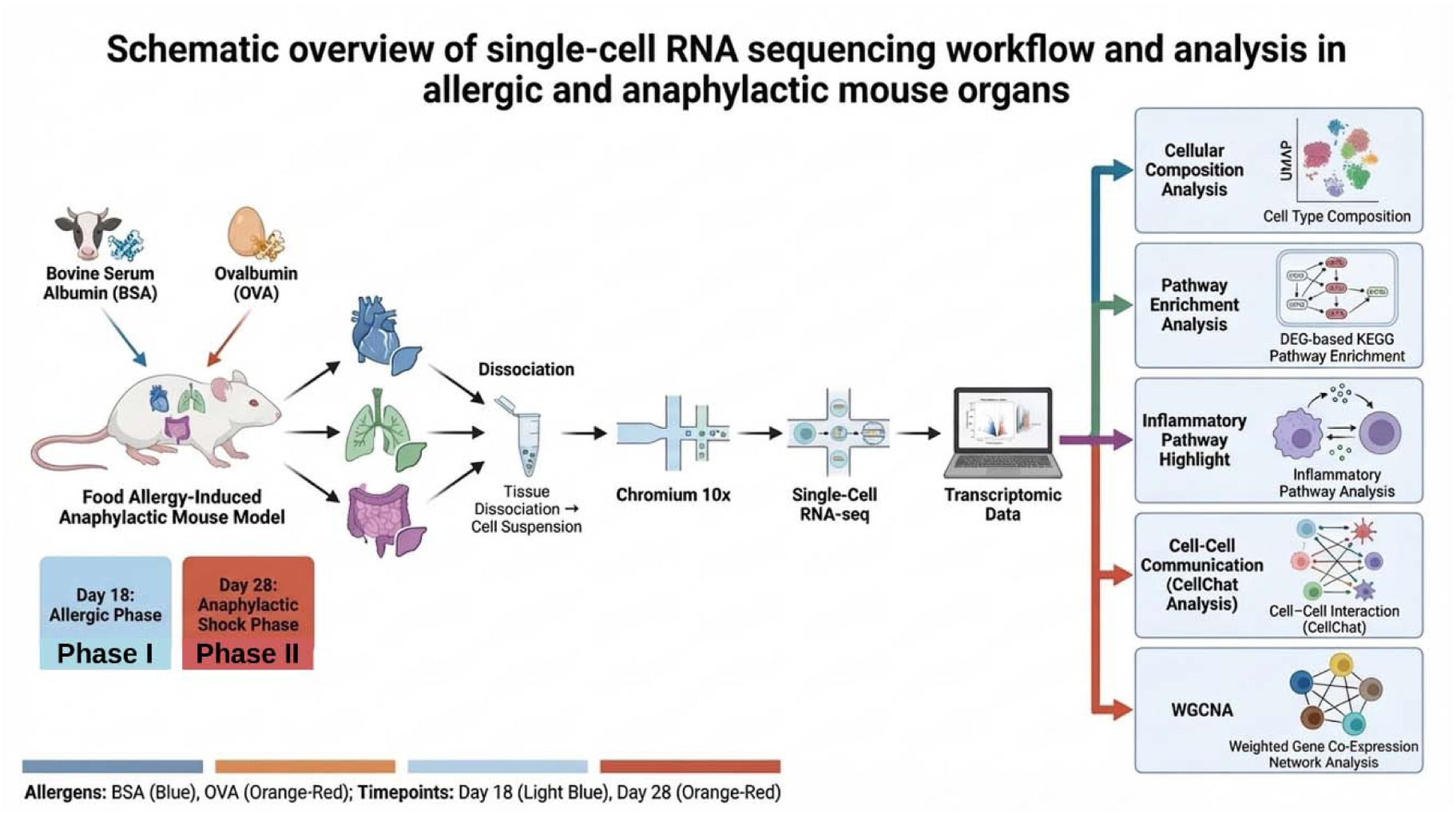

## 1. Introduction

Abnormal humoral and cellular immune responses to externally ingested antigen or stimuli results in allergic diseases such as atopic dermatitis (AD), asthma, allergic rhinitis, food allergy (FA), and at severe conditions, these pathological processes result in anaphylactic shock (AS). Due to urban lifestyles and industrialization, the incidence of allergy increased in many developed as well as developing countries with FA being the most common [1]. Exposure to small amount of food allergens might initiate mild to moderate pathological responses such as airway inflammation, gastrointestinal disorders, and urticaria. In most severe conditions, immediate hypersensitivity to food allergens causes resident and global immune/effector cell modulation, systemic proinflammatory mediator release, multi-organ failure resulting in life-threatning AS.

Although 2881 foreign proteins are documented to be allergenic (IgE mediated), only certain ones depending upon intrinsic molecular properties (structural and chemical) and systemic environmental factors elicit immediate hypersensitivity reactions to induce pro-allergic IgE or anti-allergic IgG responses [2, 3]. Dearman et al., demonstrated that ovalbumin (OVA) and bovine serum albumin (BSA), exhibited different antibody isotype (IgE/IgG) production that is reflected in discrete T lymphocyte response (Th1 and Th2) [4]. Moreover, the ratio of IgE and IgG determines the outcome of allergic response either by enhancing Th2 pro-allergic cascade or anti-allergic T-reg cascade depending upon IgE and IgG binding to FcεR1 and FcγRIIb mast cell receptors respectively [5].

Hence, the allergenicity varies based on the intrinsic biophysical and structural properties of the allergen that dictates humoral balance and clinical outcome. The most well-characterized food allergens include proteins from milk (Bos d), eggs (Gal d), peanut (Ara h), shellfish (tropomyosin) in addition to various plant based allergens [6]. However, no studies have reported comparing the magnitude of allergencity of two different allergen in the same model at single-cell resolution.

The allergic response induced by allergens is a spatio-temporally coordinated process spanning from sensitization, prodrome, early reaction, late reaction and anaphylactic reaction phases. Based upon the affected organ, each phase involves specific modulation and complex interplay between immune (dendritic cells, T and B lymphocytes, mast cells, macrophages, natural killer cells, innate lymphoid cells, eosinophils, neutrophils) and organ-specific effector cells that strictly regulate the stage-wise development of allergic pathology. These cells and their subsets release diverse mediators (antibodies, cytokines, growth factors, enzymes, alarmins, miRNAs) that can either propagate or resolve inflammation[7]. Symptom severity depends not only on IgE levels but also on cellular sensitivity, tissue-specific recruitment, and the functional programming of immune cells across organs.. Moreover, the unique susceptibility of each organ to mediators released by both resident and recruited immune cells shapes the resultant anatomical pathology.. Pucci et al. observed that bone marrow transplantation transfers systemic immune responses, whereas lung transplantation confers only asthma specific susceptibility [7]. Furthermore, aside from recipient immunoregulatory alterations, the abundance of antigen presenting cells and the cellular composition of donor liver may provoke allergic responses to previously tolerated foods [8]. Hence, understanding the temporal dynamics of the cells (inflammatory and effector) at cellular and sub-cellular level in key organs, their communication pattern and the difference in pathway modulation are crucial in understanding the intensity of immune response and resultant pathological alterations in organs during allergic to AS. Various multi-omics technologies such as genomics, transcriptomics, epigenomics, proteomics, metabolomics, exposomics, microbiomics, have been used to understand immune cells heterogeneity in food allergy [9].

Though these allergic studies have shed light on the profiling of immune cells, the key organ specific profiling of immune cells between two different food allergens at two different phase of allergy are still lacking. Three predominant organs affected during immunoglobulin mediated food allergens are cardiovascular system (plasma leak, vasodilation, myocardial depression, shock, tachycardia, hypotension), respiratory system (dyspnea, cough, wheeze, bronchoconstriction, mucus production), and digestive system (gas, bloating and diarrhea). World anaphylaxis society recently defined anaphylaxis as any compromise in breathing and/or circulation [10]. Our previous histological studies in AS murine model showed that lung and intestine showed higher infiltration of mast cells and eosinophils with enhanced expression of tryptase, receptor tyrosine kinase and induced nitric oxide synthase. Further enhanced expression of endothelial nitric oxide synthase (vasodilator) was found in heart, lung and intestine [11]. From our earlier study, we found that lungs and intestine are key organs that are affected in anaphylaxis. Mechanistically, we reported that voltage dependent potassium channels (Kv) play a significant role in hypotension and blockade of Kv channels restores blood pressure and decreases mortality rate [12, 13]. Though anaphylaxis is largely defined as an acute allergic reaction, chronic sensitization of immune cells upon allergen exposure plays a key role in the intensity of resultant anaphylaxis. Recently, American Academy of Allergy, Asthma and Immunology (AAAI) and European Academy of Allergy and Clinical Immunology (EAACI) proposed personalized or precision medicine approach to treat allergic patients. This treatment would be guided based upon the disease endotype (underlying cellular and molecular mechanisms) rather only based upon phenotype (clinical characteristics) alterations which will result in better treatment statergies [14]. Hence, in order to understand the change in higher resolution of cellular differences when exposed to two different antigens, their migration, their pathological pathways involved and their communication in specific microenvironment, at sensitization (phase I-day 18) and anaphylactic phase (phase II-day 28), we performed scRNA Seq analysis in heart, lung and intestine isolated from OVA and BSA-induced mouse model.

## 2. Results

### 2.1. Bovine Serum Albumin and Ovalbumin sensitization enhances allergic response in the experimental animals

In order to confirm that sensitization of experimental animals with BSA and OVA elicits specific immunologic IgE response, we performed Enzyme-Linked Immunosorbent Assays using plasma samples from experimental animals. We found significant increase in Total IgE (Fig 2a, *p*<0.05) and BSA specific IgE (Fig 2b, *p*<0.01) at day18 (D18) and day 28 (D28) in our BSA model. Similarly, we found significant increase in total IgE (Fig 2c, p<0.01) and OVA-specific IgE (Fig 2d) at D18 and D28 in the OVA model as well. These results demonstrate that the animals were allergic by eliciting allergen specific IgE response.

**Fig 1:**
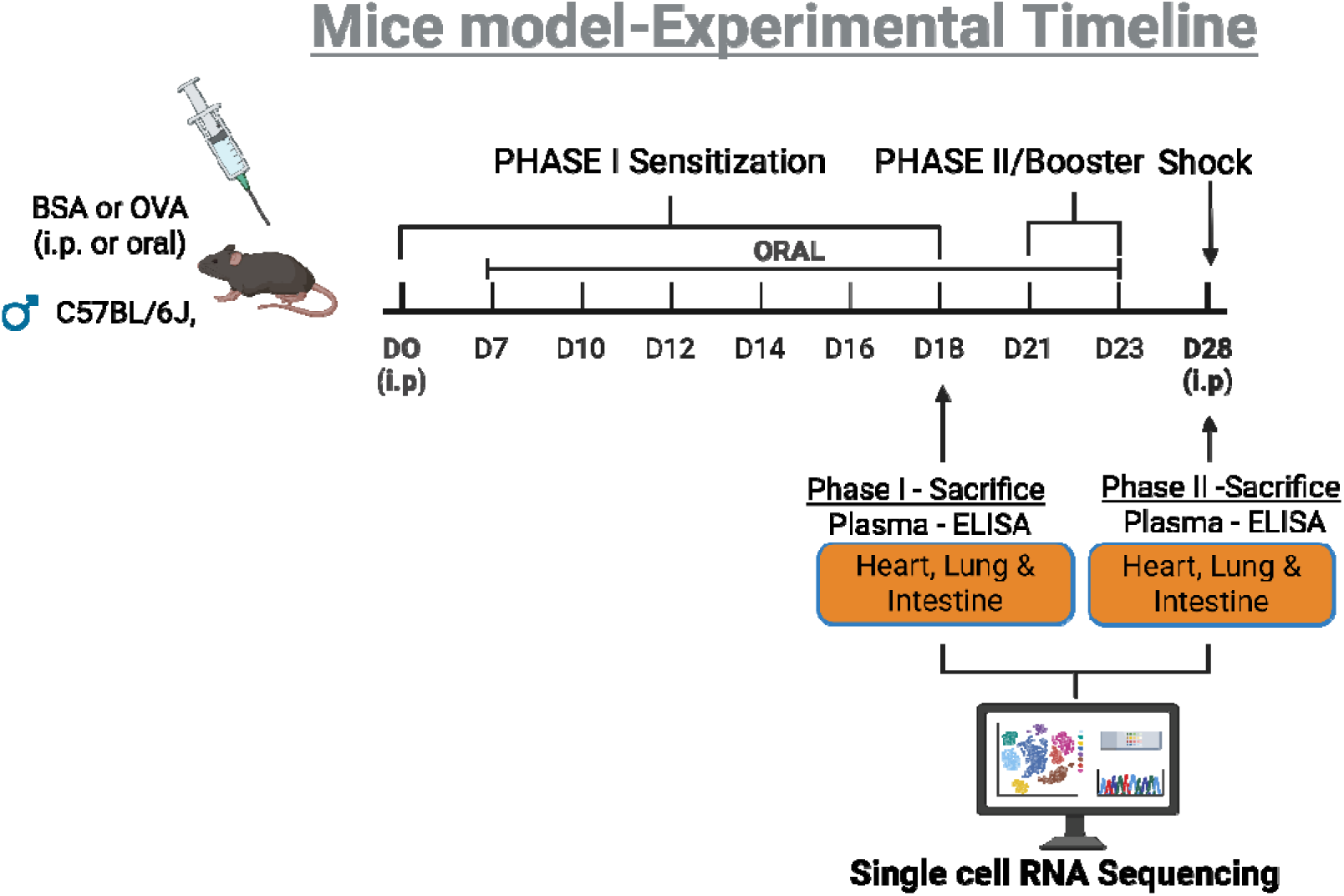
Schematic representation of sensitization schedule to create murine food allergy and anaphylactic shock model. Mice (20-25 grams) were sensitized either by intraperitoneal or oral administration of Bovine Serum Albumin (BSA) or Ovalbumin (OVA) depending upon the experimental schedule. Animals were sacrificed, blood and organs were collected on day 18 (heart, lung and intestine) and day 28 (heart, lung and intestine) for experimental analysis.

**Fig 2:**
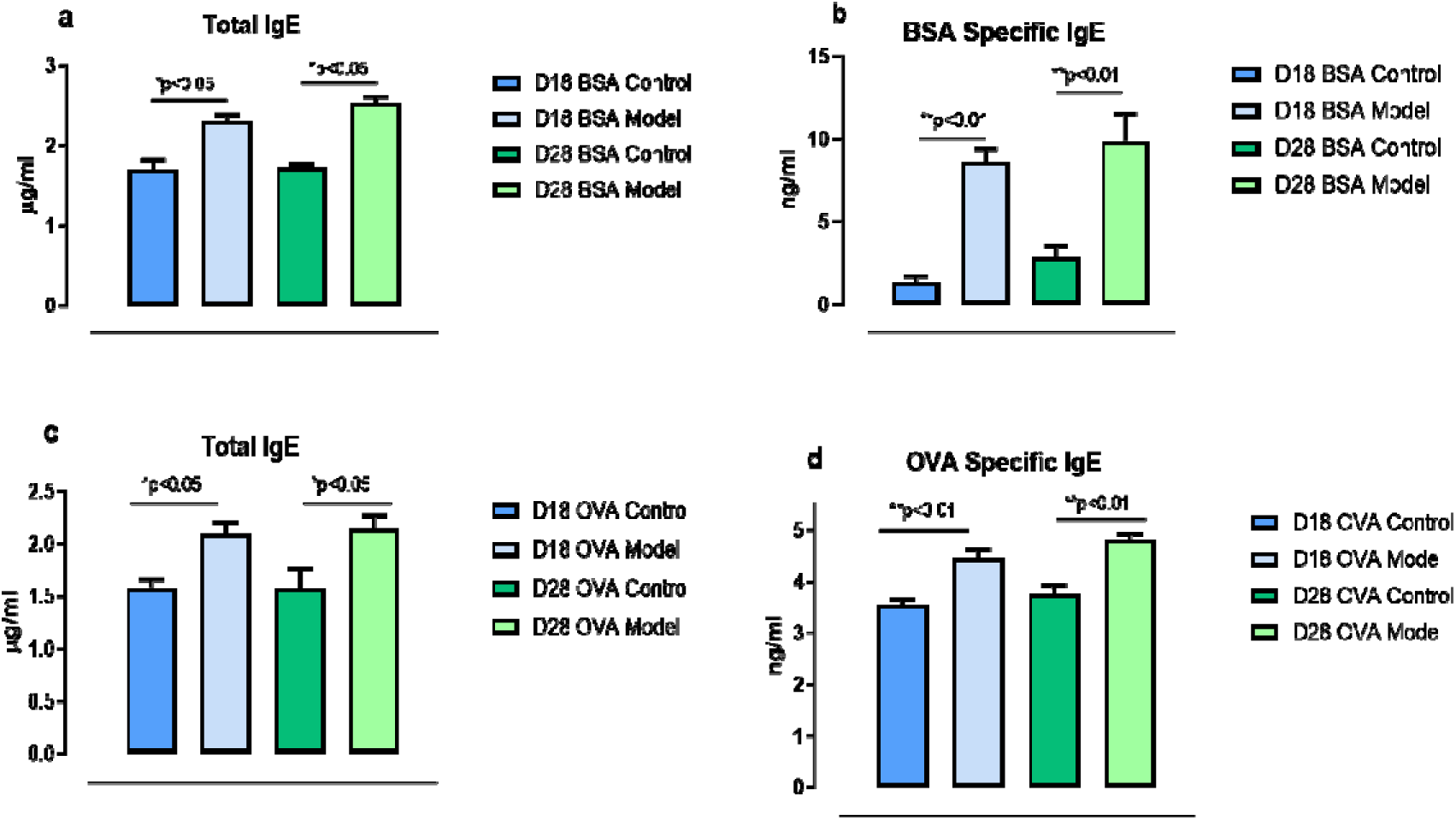
Ovalbumin sensitization enhances secretion of total IgE (OVA and BSA) and specific IgE (OVA and BSA) in the experimental animals. Serum total IgE towards BSA (a) and OVA (c) at D18 and D28 in response to allergen sensitization compared to respective vehicle controls. Total serum IgE values are represented as µg/ml. Total BSA-specific IgE (c) and OVA-specific IgE (d) at D18 and D28 in response to ovalbumin sensitization compared to respective vehicle controls. OVA and BSA-specific IgE values are represented as ng/ml. Values are represented as mean ± SEM, (n=4). *p<0.05 compared to control, **p<0.01 compared to respective controls.

### 2.2. BSA and OVA administration induced global temporal shift in cellular composition of heart, lung and intestine

In order to identify the cellular composition and their diversity in BSA and OVA models at two allergic phases, we performed unsupervised clustering of heart, lung and intestine based on the single cell sequencing data. We identified a sum of 284,191 cells in heart, 256,342 cells in lungs, and 213,024 cells in intestine. After quality filtering, the tSNE-clustering evaluation defined 14 cell type clusters in heart (Fig. 3A,3D), 16 cell clusters in lungs (Fig. 3B,3E) and 11clusters in intestine (Fig. 3C,3H). The difference in cell number at different experimental conditions of heart, lung and intestine are represented in Fig 2S1-2S6. In heart, we found significant decrease in pericytes, fibroblasts, endothelial and mesothelial cells in D18 and D28 BSA model. While in OVA model, we found significant decrease in pericytes, fibroblasts (D28) and EC (D18, D28) but mesothelial cell number increased (D18, D28). Number of certain cells such as proliferative cells, T cells, B cells were inversely correlated at D18 and D28 in BSA model. In OVA model, inverse correlation was found in proliferative cells, T cells, myeloid cells, Schwann cells and VSMC. Notably, we found significant increase in cardiomyocytes at D28 compared to D18 in both models.

**Fig 3:**
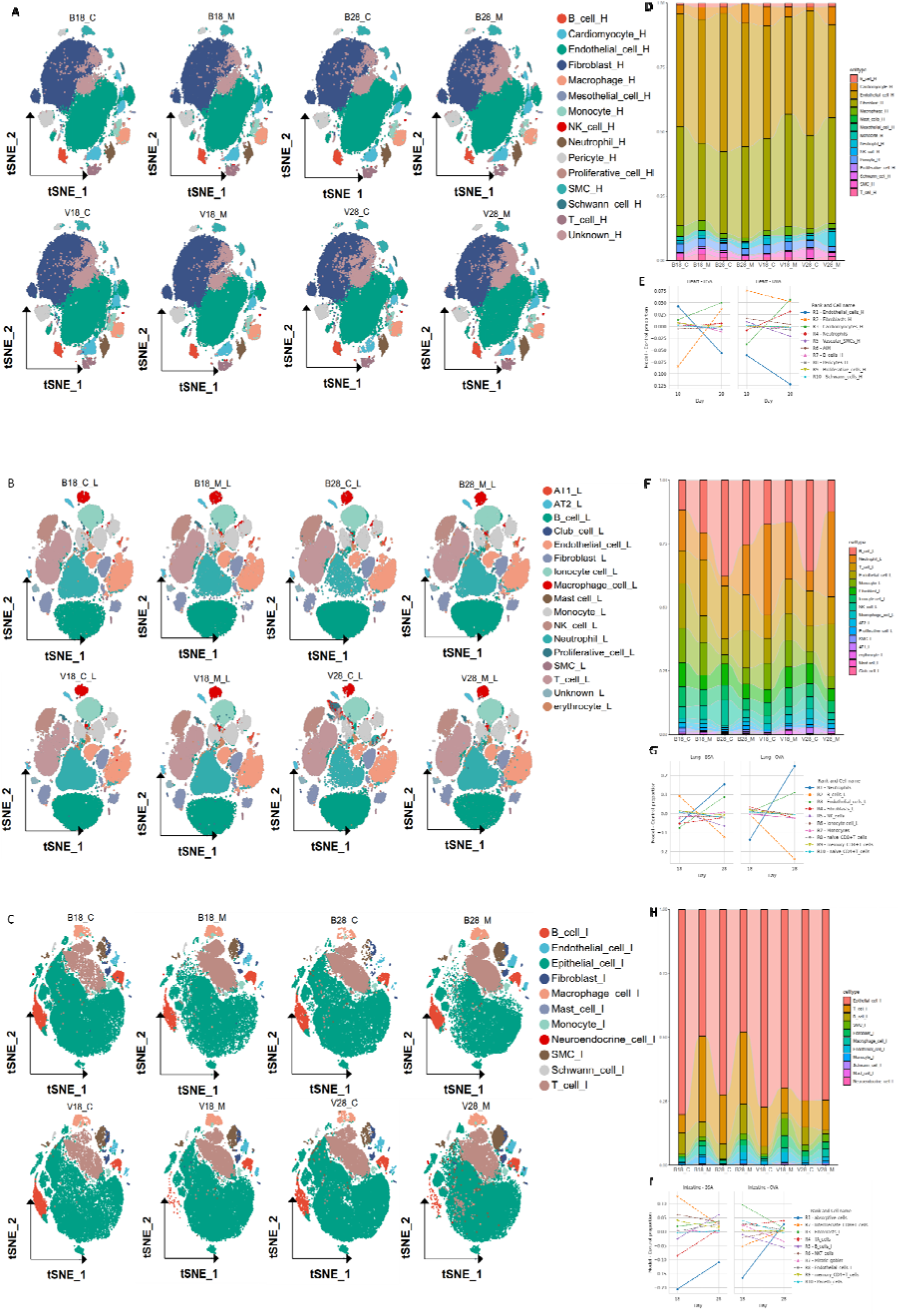
Temporal change in cellular composition of heart, lung and intestine in the experimental animals. tSNE plot of scRNA-seq data obtained after unsupervised clustering representing the changes in cellular composition of heart (A), lung (B) and intestine (C). Each dot represents a single cell and they are color coded according to the cluster identification. Bar plot depicting the changes in the cell type ratio in phase I and phase II of BSA and OVA in Heart (D), lung (F) and intestine (H). Change in top 10 cell type proportion from allergic to anaphylactic phase of heart (E), lung (G) and intestine (I) are ranked and represented as line trajectory plot.

In lungs, we found decrease in ionocyte, fibroblasts in BSA phase I and phase II compared to controls. An increase in erythrocytes and epithelial cells in both phases were observed in BSA model. Myeloid cells, ECs decreased in phase I while increased in phase II in both models. Furthermore, we found that B cells, T cells and NK cells increased in phase I and decreased in phase II. In OVA model. Almost all cells increased in phase I compared to control except myeloid cells that increased in phase II. Erythrocytes and ECs increased significantly in phase I and slightly in phase II and all other cells decreased in phase II.

During the allergic phase (day 18), intestinal sensitization to BSA and OVA elicited distinct cellular responses. The BSA model was characterized by a significant increase in T cells (in both BSA and OVA-challenged mice) and myeloid cells, alongside a reduction in epithelial cell numbers. In contrast, OVA sensitization triggered a pronounced expansion of fibroblasts and endothelial cells in both models. Notably, epithelial cell populations increased in the OVA model but decreased in the BSA model. These findings support the concept that allergen-specific patterns of intestinal immune and stromal remodeling underlie divergent pathogenic pathways in food allergy.

### 2.3. BSA and OVA modifies subcluster cell population in heart, lung and intestine

Further to identify the subclusters of cells in each organ and to evaluate the variation in more specific composition of cell types between experimental groups, we performed dimensionality reduction and cluster analysis using harmony algorithm with a resolution of 0.8 [15]. We identified 34 different subclusters in heart (Fig 4A, 4B), 31 different ones in lung (Fig 4F, 4G), and 43 ones in intestine (Fig 4K, 4M). Marker gene annotations identified 11 types of T cells and 12 different myeloid cells in heart, 12 types of T cells and 12 different myeloid cells in lungs, 12 types of T cells, 12 different myeloid cells and 11 different epithelial cells in the intestine.

**Fig 4:**
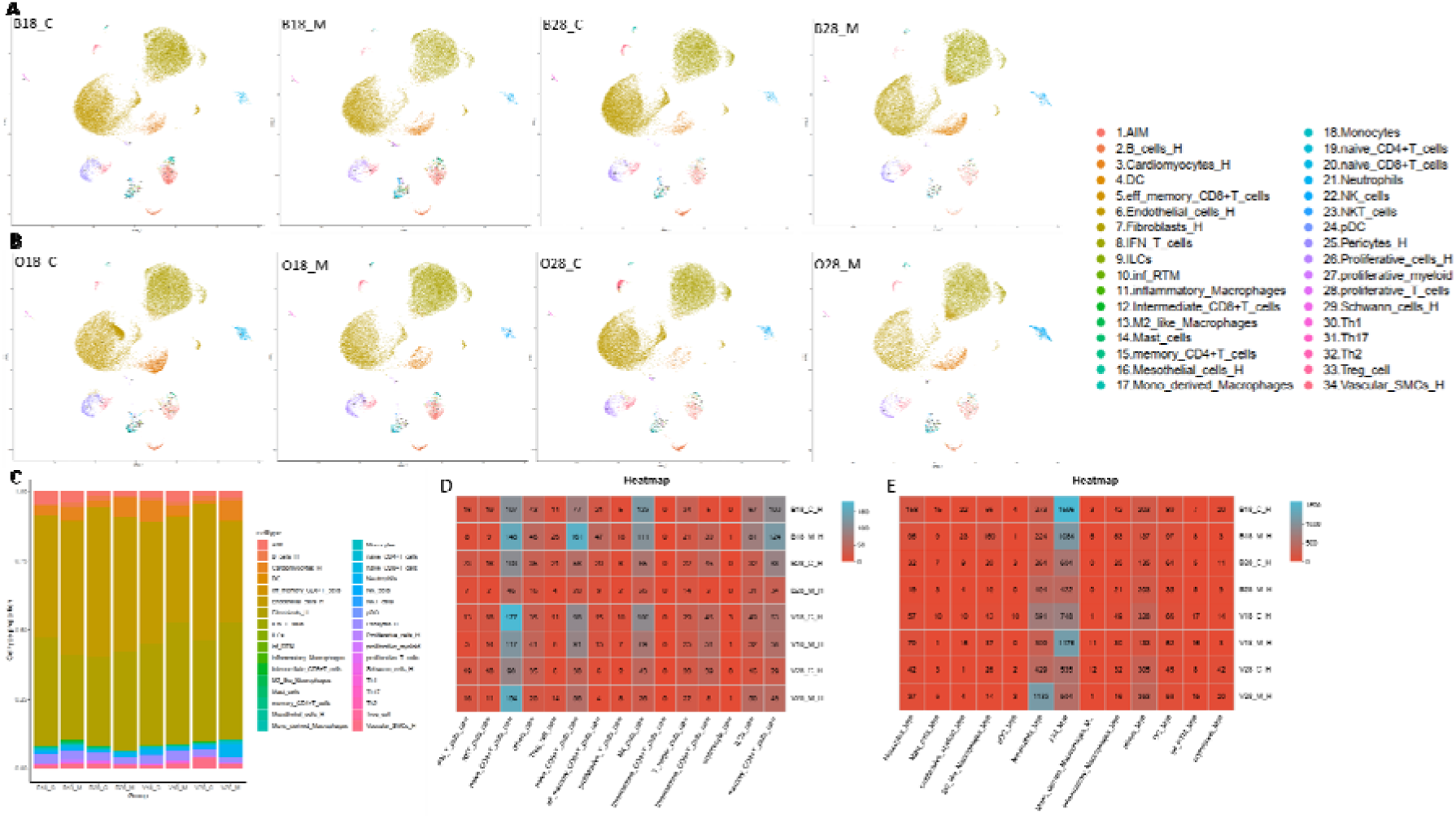
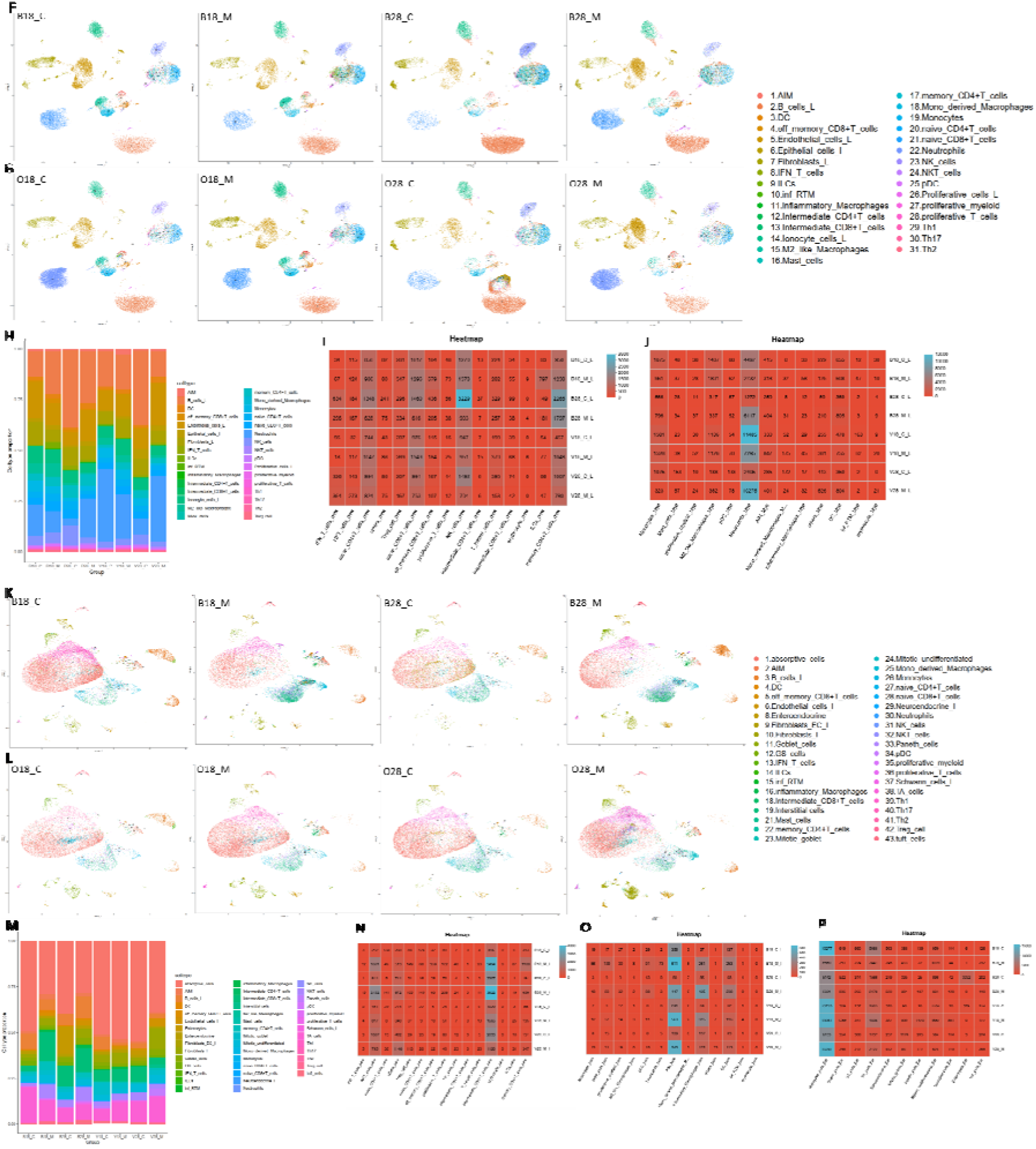
Representation of global sub-cellular population of heart, lung and intestine in the experimental samples. UMAP representation of change in temporal dynamics of subclusters in BSA and OVA model of heart (A, B), lung (F, G) and intestine (K, L). Bar plot representation depicting the change in subcluster proportion in phase I and phase II in BSA and OVA model of heart (C), lung (H), and intestine (M). Heatmap representation of change in cell number in T cell subcluster (D, I, N), myeloid cell subcluster (E, J, O) and epithelial cell subcluster (P) in heart, lung and intestine respectively. Y-axis in the heatmap represents BSA (B) or OVA (O), phase I (18) or phase II (28) and control (c) or model (M). X axis represent the subcluster type (T cell, myeloid cell or epithelial cell).

In heart, T-cell signature reveals opposite patterns in cell number at D18 and D28 in naïve CD8+ T cell, T helper cells, ILCs and memory CD4+ T cells between BSA and OVA group (Fig 4D). Analysis of myeloid cells showed significant increased in neutrophils at D28 compared to D18 in both BSA and OVA models (Fig 4E). Erythrocytes were decreased in both models in both phases, The number of monocytes decreased in both phases in BSA model but in OVA model, the number of monocytes increased at D18 and decreased in D28. Mast cell, an important player in anaphylaxis, decreased in BSA model but increased in phase II in OVA model.

In lung, we found decrease in IFN-T cells in phase I and II in BSA model. However, in OVA model, the number of cells decreased in phase I but increased in phase II. Naïve CD4+ T cell, Naïve CD8+ T cell, memory effector CD8+ T cell, intermediate CD8+ T cell and memory CD4+ T cells increased in phase I and decreased in phase II in both models. NKT and NK cells increased in phase I and decreased in phase II in BSA model. However, in OVA model, we found increase in NKT cell and decrease in NK cells in phase II. Similar trend was observed in Tregs. Myeloid subcluster analysis showed that neutrophils decrease in phase I and increased in phase II in both models. Mast cell increased in phase I in both models. In OVA model, mast cell decreased in phase II. Similar trend was found in proliferative myeloid cells and M2 like macrophages increased in both phases in both models.

In the intestine, the number of IFN-T cells, Tregs, effector memory CD+8 T cells, NK, ILC, memory CD4+ T cell were increased in both phases in both models. However, NKT cells, proliferative cells, intermediate CD8+ T cells were increased in phase I in BSA model but increased in phase I and decreased in phase II of OVA model (Fig 4N). In myeloid cell population, monocytes, mast cells, plasmacytoid dendritic cells, dendritic cells increased in phase I and phase II of both models. Neutrophils, increased in phase I but decreased in phase II in both models which was opposite to lungs. Inflammatory macrophages increased in phase I and phase II of BSA model. In OVA model, the inflammatory macrophages increased in phase I but decreased in phase II (Fig 4O). Additionally, epithelial population of intestine showed most of the cell number decreased in phase I (absorptive, goblet, GS, TA, enteroendocrine, paneth, interstitial, enterocytes) and increased in phase II in BSA model, while we found the opposite trend in the same cell populations in OVA model(Fig 4P).

### 2.4. Cell Specific pathway enrichment and trajectory analysis across multiple organs

Differential gene expression (DEG) analysis was performed to identify the top 200 DEG in each organ-allergen-phase combination in each major cell type. Pathway enrichment analysis was performed to determine the top 5 inflammatory and global KEGG pathways and their trajectory analysis was conducted to monitor temporal dynamics of selected pathways from allergic (phase I) to shock phase (phase II) based on mean_log2FC values. To map and annotate this cellular atlas deeply, we also identified certain pathways that were modulated in allergic phase in certain cells and the same pathway modulated in other specific cells at shock phase (Fig 5).

**Fig 5:**
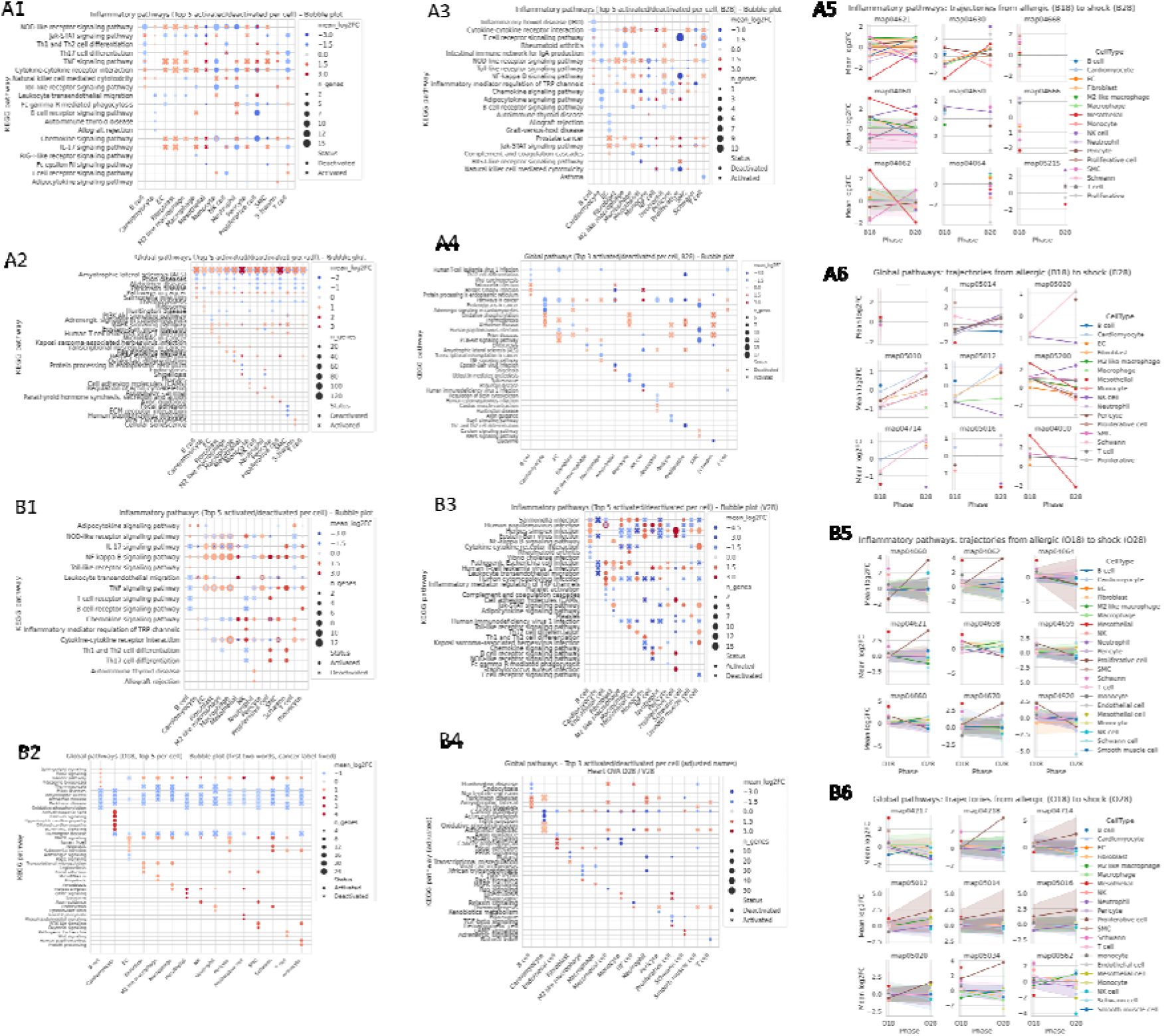
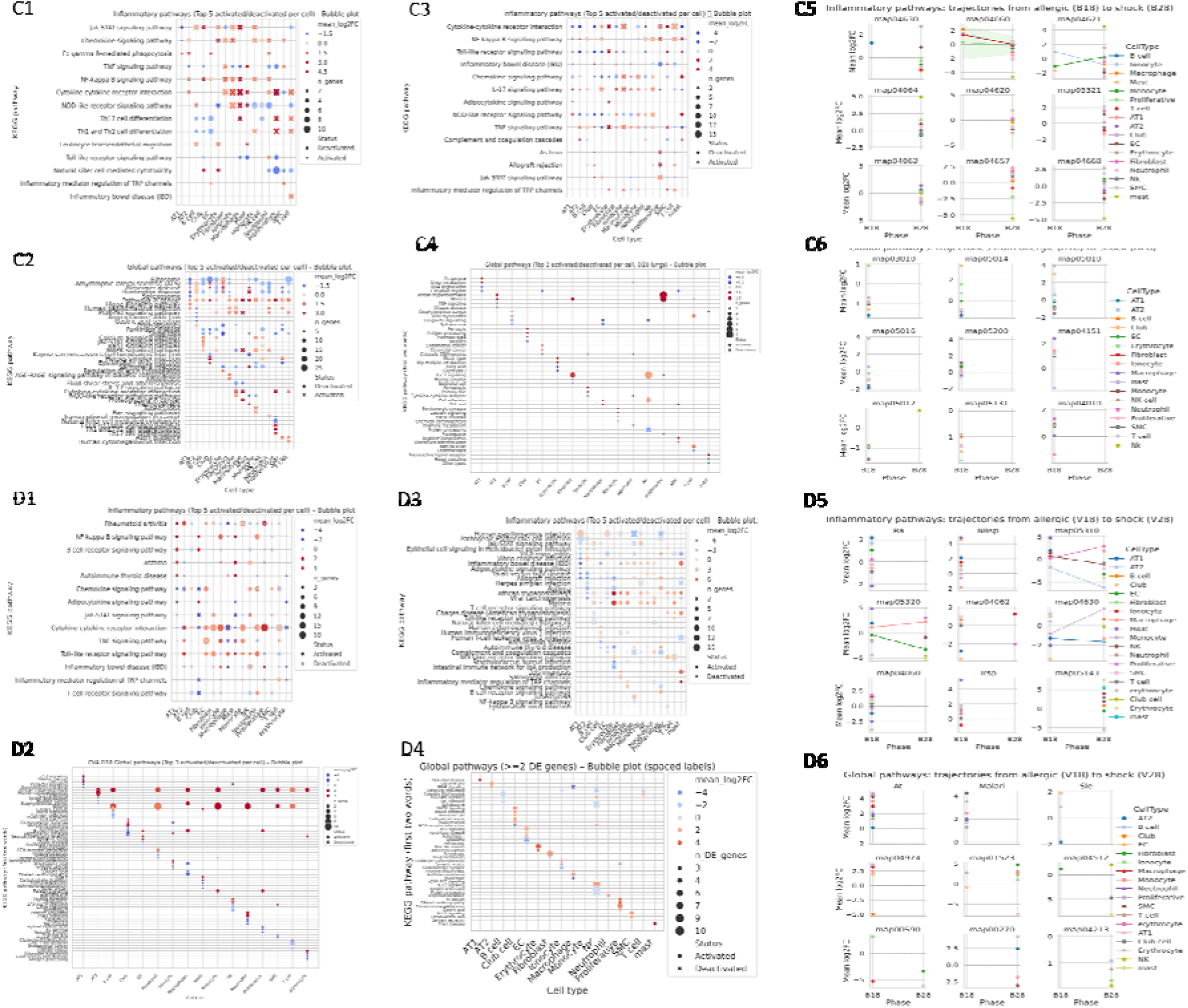
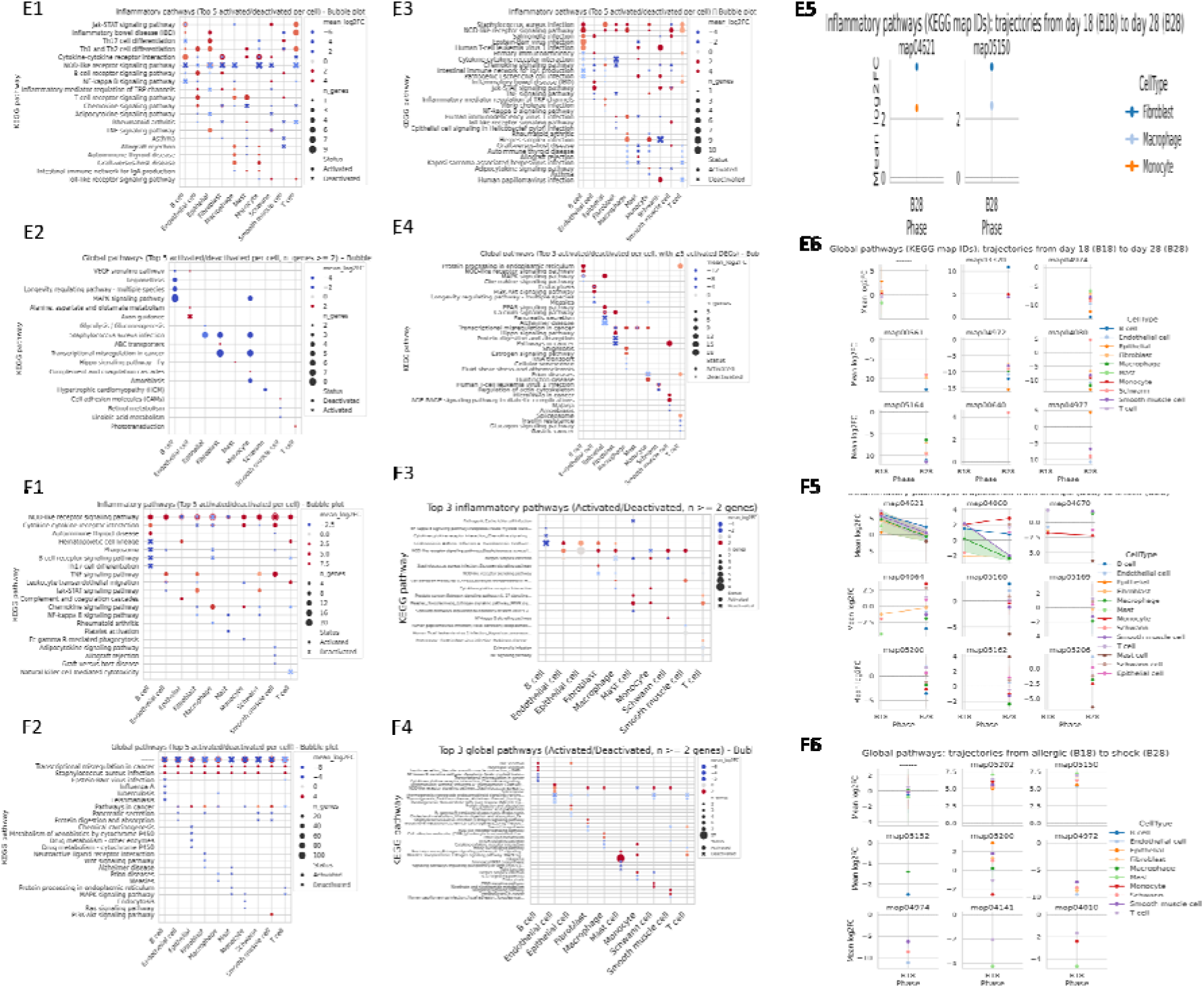
Multi-organ pathway analysis and temporal trajectory analysis in experimental animals. Top 200 DEG’s in each cell was used to analyze pathway enrichment and identify top activated/deactivated inflammatory and global pathways. Cardiac (A,B) transcriptional responses across 15 cell types. Bubble-map representation of inflammatory (A1,A3,B1,B3) and global (A2,A4,B2,B4) KEGG pathway enrichment at D18 and D28 for BSA and OVA models respectively. Temporal trajectory plots of top 5 inflammatory (A5, B5) pathways and global (A6,B6) pathways showing cell-type-specific (log2FC) progression from D18 to D28 in BSA and OVA models respectively. Pulmonary transcriptional responses across 16 cell types. Bubble-map representation of inflammatory (C1,C3,D1,D3) and global (C2,C4,D2,D4) KEGG pathway enrichment at D18 and D28 for BSA and OVA models respectively. Temporal trajectory plots of top 5 inflammatory (C5,D5) pathways and global (C6,D6) pathways showing cell-type-specific (log2FC) progression from D18 to D28 in BSA and OVA models respectively. Intestinal transcriptional responses across 10 cell types. Bubble-map representation of inflammatory and global KEGG pathway enrichment at D18 and D28 for BSA and OVA models. Heatmap representation of inflammatory (E1,E3,F1,F3) and global (E2,E4,F2,F4) KEGG pathway enrichment at D18 and D28 for BSA and OVA models respectively. Temporal trajectory plots of top 5 inflammatory (E5, F5) pathways and global (E6,F6) pathways showing cell-type-specific (log2FC) progression from D18 to D28 in BSA and OVA models respectively. Bubble plot-Color represents mean_log2FC score and n represents number of DEG’s in specific pathway. Pathway trajectory analysis-each color represents a cell type and the KEGG mapid of corresponding pathways are mentioned in facet above each pathway trajectory representation. Values are represented as mean_log2FC.

In the heart, 15 different cell types were identified by scRNA-seq analysis for both models of allergen. Following BSA challenge, 5,776 DEGs were identified at D18 and 5,292 DEGs at D28. Pathway enrichment at D18 showed activation of important inflammatory pathways such as NOD-like receptor signaling pathway (map04621), cytokine-cytokine receptor interaction (map04060) and chemokine signaling pathway (map04062) (Fig 5A1). By D28 the inflammatory response was enhanced by enrichment of 21 different inflammatory pathways, mainly NOD-like receptor signaling pathway (map04621), NF-kappa B signaling pathway (NF-κB, map04064), JAK-STAT pathway (map04630). In the OVA model, 5,276 DEGs were found at D18, which increased to 5,692 DEGs at D28. OVA-induced pathway enrichment at D18 shows enrichment of cytokine-cytokine receptor interaction (map04060), IL-17 signaling pathway (map04657) and TNF-α signaling pathway (map04668). At D28, 30 inflammatory pathways were enriched, comprising platelet activation, JAK-STAT and NF-κB pathways as well as 35 top global pathways including pathways related to thermogenesis, focal adhesion etc. Pathway trajectory showed temporal dynamics of the top 5 inflammatory and global pathways, representing the progression of early allergic responses to severe anaphylactic reactions in heart tissue. Inflammatory pathways include cytokine-cytokine receptor interaction, chemokine signaling, leukocyte trans endothelial migration.

In the lung, the scRNA-seq analysis defined 16 different cell types in both allergen models. Following BSA challenge, 5,966 DEGs were detected at D18 progressing to 6,041 DEGs at D28. Pathway enrichment at D18 showed that inflammatory pathways such as NOD-like receptor signaling pathway (map04621, the most prominent with 26 occurrences), chemokine signaling pathway (map04062), TNF-α signaling pathway (map04668) and Th17 cell differentiation (map04659) were activated. In day 28, in addition to NF-κB, Toll-like receptor, IL-17, pathways related to asthma, inflammatory regulator of TRP channels are modulated as well.

By D28, global response grew stronger with enrichment of more pathways such as african trypanosomiasis (map05143) and graft-versus-host disease (map05332) related pathways, calcium signaling, etc. In the OVA model, 5,999 DEGs were detected at D18 and the number of DEGs increased to 6,203 at D28. OVA-induced pathway enrichment at D18 revealed cytokine-cytokine receptor interaction (map04060; 22 occurrences), rheumatoid arthritis (map05323; 20 occurrences), TNF-α signaling pathway (map04668), toll-like receptor signaling pathway (map04620) and chemokine signaling pathway (map04062). At D28, 36 inflammatory pathways were enriched, including those related to african trypanosomiasis (map05143), asthma (map05310), allograft rejection (map05330), inflammatory bowel disease/IBD (map05321), and malaria (map05144) in addition to 68 global pathways (top 3). Pathway trajectory analysis showed temporal dynamics of top 5 inflammatory and global pathways depicting the progression from early allergic response to severe anaphylactic response in lung tissue.

In the intestine, scRNA-seq analysis revealed the existence of 10 different cell types for both of the allergen models. Following BSA challenge, 3,962 DEGs were identified at D18 which enhanced to 3,644 DEGs at D28. Pathway enrichment on D18 showed activation of major inflammatory pathways, such as NOD-like receptor signaling pathway (map04621), cytokine-cytokine receptor interaction (map04060) and Th1 and Th2 cell differentiation (map04658). By D28 the inflammatory reaction showed increasing strength with enrichment of 28 different inflammatory pathways, mostly chemokine signaling pathway (map04062), NOD-like receptor, JAK-STAT and staphylococcus aureus infection (map05150)-related pathways. In the OVA model, a total of 3,778 DEGs were found at D18 that increased to 4,000 DEGs at D28. OVA-induced pathway enrichment at D18 which highlighted NOD-like receptor signaling pathway (map04621, the most prominent pathway, 39 occurrences), cytokine-cytokine receptor interaction (map04060), and leukocyte transendothelial migration (map04670). At D28, 75 inflammatory pathways were enriched, hence top 3 inflammatory pathways in which 2 or more genes (activated/deactivated) in specific pathways are represented, this include cell adhesion molecules/CAMs (map04514), leukocyte transendothelial migration (map04670), tight junction (map04530), pathogenic Escherichia coli infection (map05130) and a comprehensive suite of immune related pathways including asthma (map05310), natural killer cell-mediated cytotoxicity (map04650), Fc epsilon RI signaling pathway (map04664), intestinal immune network for IgA production (map04672), and inflammatory bowel disease/IBD. Pathway Trajectory analysis showed the temporal dynamics of the top 5 inflammatory and global pathways showing the progression of early allergic responses to severe anaphylactic responses in intestine tissue.

Cross-organ comparison found both common and organ specific transcriptional responses to food allergen challenge. Key common inflammatory pathways between all three organs were NOD-like receptor signaling (map04621), Jak-STAT signaling (map04630), cytokine-cytokine receptor interaction (map04060), TNF-α signaling (map04668) and NF-kappa B signaling (map04064) pathways. The kinetics and scale of pathway activation was substantially different across organs.

Pathway trajectory examination in both BSA and OVA models identified the conservation of inflammatory mechanisms underlying food allergy-induced anaphylaxis, in addition to identifying allergen-specific transcriptional signatures in each organ. Notably, both allergens induced signaling pathways associated with NOD-like receptors (map04621) and cytokines and cytokine receptors (map04060) in all organs suggesting the presence of common innate immune recognition mechanisms. OVA allergen resulted in greater diversity and increased numbers of enriched pathways at D28 compared to BSA and may emphasize differential allergen immunogenicity or more severe anaphylactic responses.

### 2.5. Intercellular communication analysis in heart, lung and intestine

To supplement this cell-autonomic view, we conducted CellChat analysis to decode intercellular networks of communication systematically since AS comprises coordinated multi-cellular reactions (Fig 6). In heart, lung and intestine, the allergic phase (D18) resulted in an extensive stromal-immune interaction with prominent antigen-presentation and adhesion modules, whereas AS (D28) resulted in neutrophil-vascular crosstalk and danger-associated signaling. We observed that the number of inferred interactions and their strength in the heart were higher at phase I in both BSA (5240;103.94) and OVA (5158;80.52) group compared to BSA (3488;67.13) and OVA (3913;76.86) in phase II. D18 networks featured in the heart were overtaken by a broad immune targeting through an APP–CD74 signaling by a variety of compartments to B cells and myeloid populations (APP_CD74) in line with increased antigen-processing communication. In Heart D18 we observed strong cytotoxic-immune recruitment with NK-to-neutrophil chemotaxis (CCL5_CCR1), alongside macrophage-inhibitory/identity signaling toward myeloid targets (PTPRC_MRC1). Nevertheless, paradoxically, neutrophil–endothelial communication became more robust, and thrombospondin-mediated interactions between the endothelium and the vascular system increased in the leading roles (THBS1_CD36) meaning that the endothelium became engaged in the context of the systemic collapse. Simultaneously, the D18-D28 collapse of mast cell-binding to extracellular-matrix scaffolds in heart attenuation was observed during multiple extracellular-matrix anchoring interactions with mast cells to laminin/fibronectin to CD44 axes (COL1A1_CD44, COL1A2_CD44, COL4A1_CD44, LAMA2_CD44, FN1_CD44) which suggested phase-dependent remodelling of mast-cell niche interactions. The neutrophil thrombospondin programs (THBS1_CD36, THBS1_CD47, THBS1_SDC4) in OVA-induced heart shock exhibited particularly significant increases, suggesting that neutrophilia therapeutic checkpoints (THBS1, THBS1, THBS1) were neutrophil-dependent.

**Fig 6:**
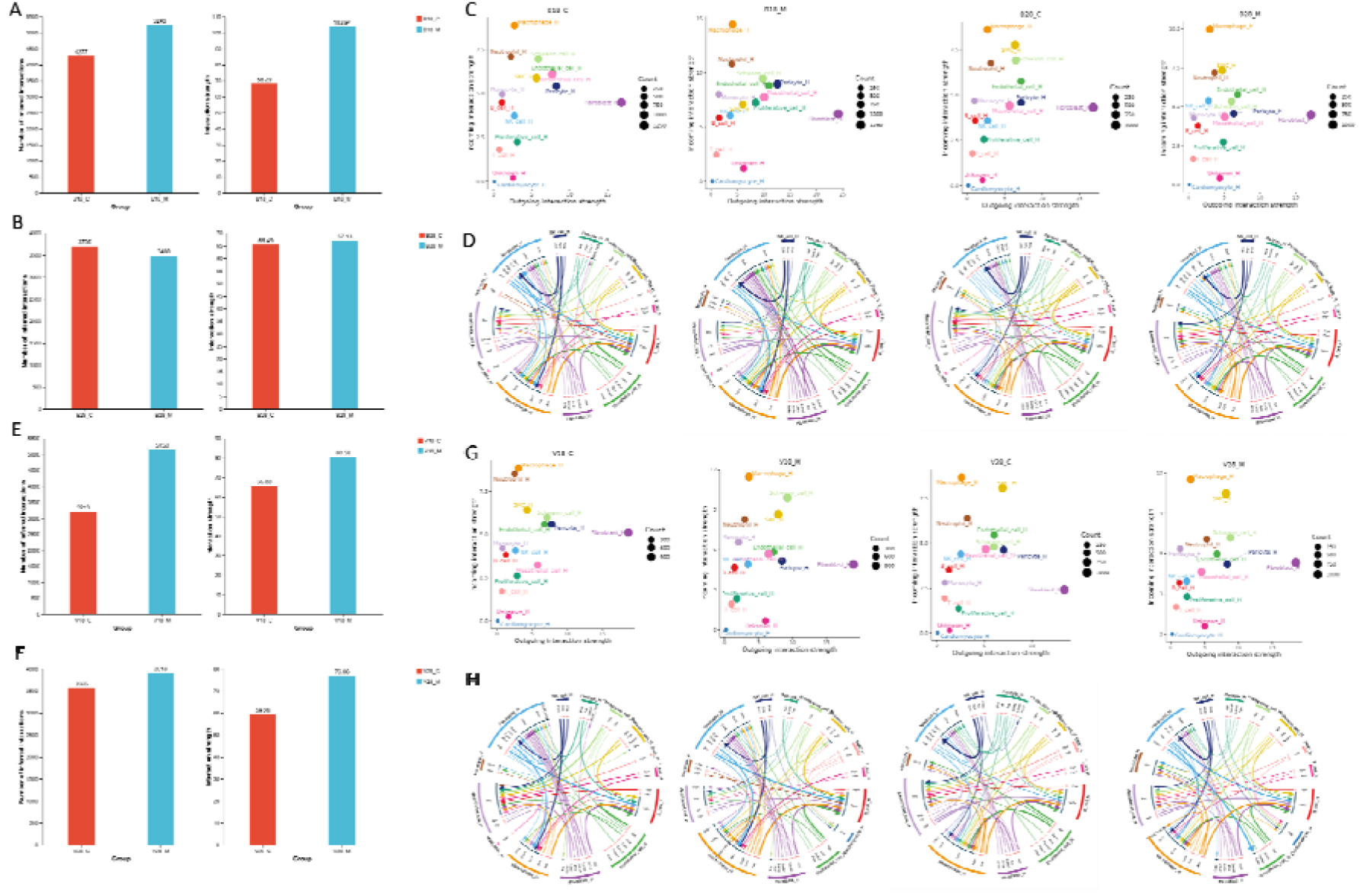
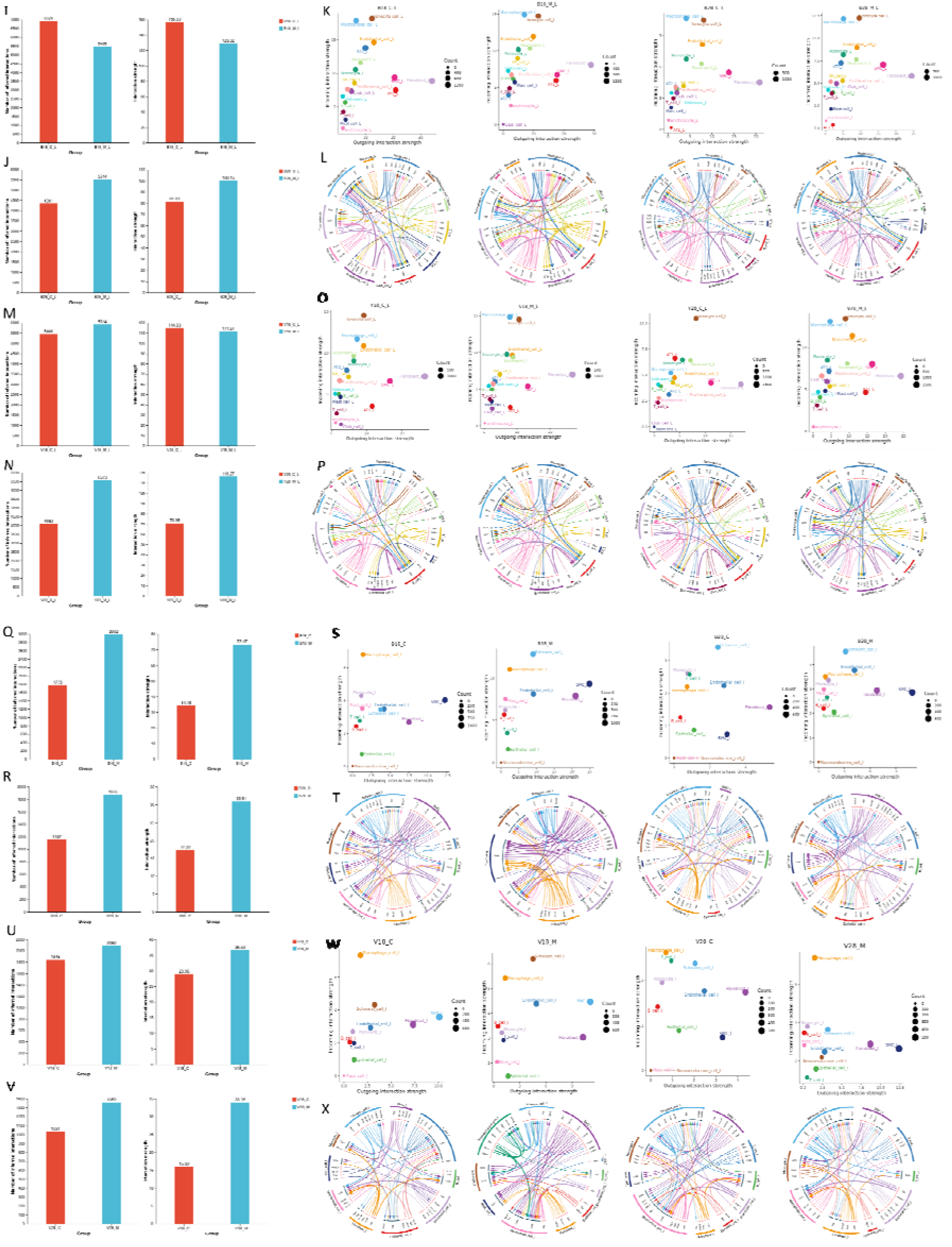
Cell chat analysis illustrating intercellular communication strength in heart, lung and intestine in different experimental conditions. The total number of inferred interactions and the interaction strength between the cells in heart (A, B, E, F), lung (I, J, M, N) and intestine (Q, R, U, V). 2D space plot representation of outgoing and incoming interaction strength between the cells at different experimental conditions in heart (C, G), lung (K, O) and intestine (S, W). Facet label above 2D plot represents BSA (B) or OVA (O), phase I (18) or phase II (28) and control (c) or model (M). iTALK plot representing top 50 ligand-receptor interaction between cells in heart (D, H), lung (L, P) and intestine (T, X) in BSA and OVA experimental groups. The outer layer represents the cell type and the inner layer represents either the ligand sent or the receptor receiving the signals. The connections were indicated in different colors representing the cell type and the thickness of the connection represents the strength of the interaction signal.

Alternatively in the lungs, the number of inferred interactions and their strength have decreased in phase I BSA (5,485;129.32) and slightly increased in OVA phase I (5914;111.51). However, these interactions were increased in phase II BSA (5514;100.15) and OVA (5673;116.27). The interaction landscape of the lung was on the whole more interaction-rich than intestine and heart, and highly enriched with leukocyte-recruiting and leukocyte-vascular coupling interactions like Neutrophil-Macrophage (APP-CD74) and Ionocyte-B_cell (APP-CD74). In line with a barrier tissue that is rich in recruitment, D18 lung also exhibited a most striking cytotoxic-immune recruitment axis NK-Neutrophil (Ccl55Ccr1) as one of the most likely edges.

We also evaluated the pattern in the intestine and as expected irrelevant on the phases, both phase I of BSA (2,992;73.47) and OVA (2,090;36.63) and phase II of BSA (1,876;30.91) and OVA (1,359;33.98) had higher number of inferred interactions and interaction strength compared to controls (Fig 6Q, R, U, V). The mixed program of myeloid cell vascular/stromal instruction, mast-cell niche anchoring and robust neuro-associated wiring were found in the intestine D18 networks. The most likely edges were Endothelia-Macrophage (APP-CD74) and Fibroblast-Macrophage (APP-CD74), which are in accordance with the widespread antigen-processing signaling to myeloid compartments. Simultaneously, the intestine demonstrated a significant ECM-mediated niche communication into mast cells, which is mediated by SMC-Mast (COLLA1-CD44) and SMC-Mast (COLLA2–CD44), neuro/glial and neuroendocrine self-communication such as Schwann-Schwann (NCAM1–NCAM1) and Neuroendocrine-Neuroendocrine (CADM1–CADM1; NRXN1–NLGN1). In the case of intestinal shock (D28), the network changed to even more dominant dominance of the APP-CD74 axis in all compartments, top edges dominated by Endothelial-Macrophage (APP-CD74), Endothelial-Monocyte (APP-CD74), Schwann-Macrophage (APP-CD74), and Fibroblast-Macrophage (APP-CD74). Incidentally, D18 top and D28 top rankings show a relative shift in favor of broad antigen-processing and myeloid-directed communication (including Macrophage-Macrophage (APP-CD74).

Heart was the only organ showing the most pronounced switch of stromal-anchoring contacts to neutrophil-endothelial signaling (THBS1-CD36) upon shock. In comparison, lung communication was consistently and persistently elevated and inflammatory in all phases. Intestine is more likely to transition from priming-to-acute as anticipated by barrier-organ initiation at D18 then systemic reaction at D28. These data thus suggest allergic priming is maintained by stromal and antigen-presentation communication (APP-CD74) and shock where neutrophils–vascular communication (THBS1-CD36) is increased and directed by organ-specific interaction strength and direction. Detailed signal flow commuication pattern between all the cells in all the organs have been provided in the supplementary figure (Fig 6S).

### 2.6. Allergen exposure alters gene co-expression network that are phase and tissue dependent

Following characterization of cellular pathway using KEGG pathway enrichment and cell chat analysis to understand intercellular communication, we employed WGCNA to identify co-expressed gene network that collectively captures biological process which is not accessible in individual pathway analysis. In heart (Fig 7B), immune activation related analysis showed that the model group had higher ORs for “cellular response to interleukin-1” in endothelial cells (model: 5.74 vs. control: 1.48) and fibroblasts (model: 5.71 vs. control: 0.99), indicating an increase in inflammatory pathways. T cells in the model had a considerably higher “T cell receptor signaling pathway” (OR = 5.28) than the control group (OR = NA, indicating minimum activity). In comparison to control cells (OR = 3.66), NK cells showed higher enrichment for “stimulatory C-type lectin receptor signaling” in the model (OR = 4.67). Cardiomyocyte function related analysis showed that it was marginally lower in the model (OR = 4.27) than in the control (OR = 5.95), “smooth muscle cell differentiation” was nonetheless high in SMCs in both groups. Compared to the control (OR = 0.99), cardiomyocyte-specific modules such as “regulation of heart contraction” were more enriched in the model (OR = 4.26), indicating compensatory adaptation. Higher recruitment was shown by neutrophils’ higher ORs for “neutrophil chemotaxis” in the model (OR = 2.87) as compared to the control (OR = 1.60). Compared with the control group (OR = 1.30), macrophages exhibited higher “antigen processing and presentation” in the model. However, in phase II we observed enhanced inflammatory response, in comparison to the control (OR = 2.36), “neutrophil chemotaxis” was much higher in the model (OR = 4.67 in neutrophils). The model showed significantly more mast cells degranulation (OR = 6.21) than the control group (OR = 7.15, but with a lower cell representation). In contrast to the control (OR = 0.62 and 0.33), fibroblasts and endothelial cells exhibited greater enrichment for “positive regulation of endothelial cell proliferation” in the model (OR = 4.59 and 5.00, respectively), indicating active vascular healing. While the level of “extracellular matrix organization” in fibroblasts was nonetheless high in both groups, it was higher in the control group (OR = 7.37) than in the model group (OR = NA, data lacking). Compared to the control (OR = 4.75 in proliferative cells), the “T cell receptor signaling pathway” was more active in the model’s T cells. In contrast to the control (OR = 3.60), B cell differentiation was higher in the model (OR = 4.21). In the OVA model (Fig 6D), indicating increased T cell activation, T cell receptor signaling was significantly higher in the model’s T cells (OR = 5.82) than in the control group (OR = 4.61). Compared to the control (OR = 3.86 for B cell receptor signaling), the model’s B cell activation was higher (OR = 4.99 in B cells). There may be structural changes in disease, since smooth muscle cell differentiation was more noticeable in the control group’s SMCs (OR = 6.72) compared to the model group (OR = 5.11 for extracellular matrix organization in SMCs). Cardiomyocytes in the model had increased sarcomere structure, which is essential for heart function (OR = 2.32), but the control group did not have it (OR = NA). Angiogenesis and vascular response processes showed that in contrast to control (OR = 4.48 for heart development), angiogenesis was significantly more prevalent in endothelial cells in the model (OR = 5.92), indicating adaptive vascular alterations. In the model, fibroblasts (OR = 1.50) and endothelial cells (OR = 1.44) had greater ORs for blood pressure regulation than the control group (OR = 1.27 and 1.81, respectively). In phase II model, immune cell recruitment and activities showed that control revealed decreased ORs (e.g., OR = 2.82 in monocytes), neutrophil chemotaxis remained higher in the model, particularly in neutrophils (OR = 3.50) and monocytes (OR = 2.96). In the model, macrophages presented antigens via MHC class II at a much higher rate (OR = 4.75) than the control group (OR = 3.01 in B cells). Cardiac and structural adaptation analysis showed that disease-driven remodeling was shown by the model’s significantly higher smooth muscle cell differentiation (OR = 6.29) compared to the control group (OR = 1.89). Prolonged cardiac stress responses were indicated by the greater presence of sarcomere organization in the model’s cardiomyocytes (OR = 4.45) compared to the control group (OR = 4.48). Metabolic responses showed that hypoxia was linked to the course of the disease because the model’s SMCs responded more strongly to oxygen levels (OR = 2.88) than the control group (OR = 3.02 in SMCs). Both groups’ fibroblasts had consistently high levels of extracellular matrix organization (model: NA; control: 4.12), while the control group’s engagement of cell types was more extensive.

**Fig 7:**
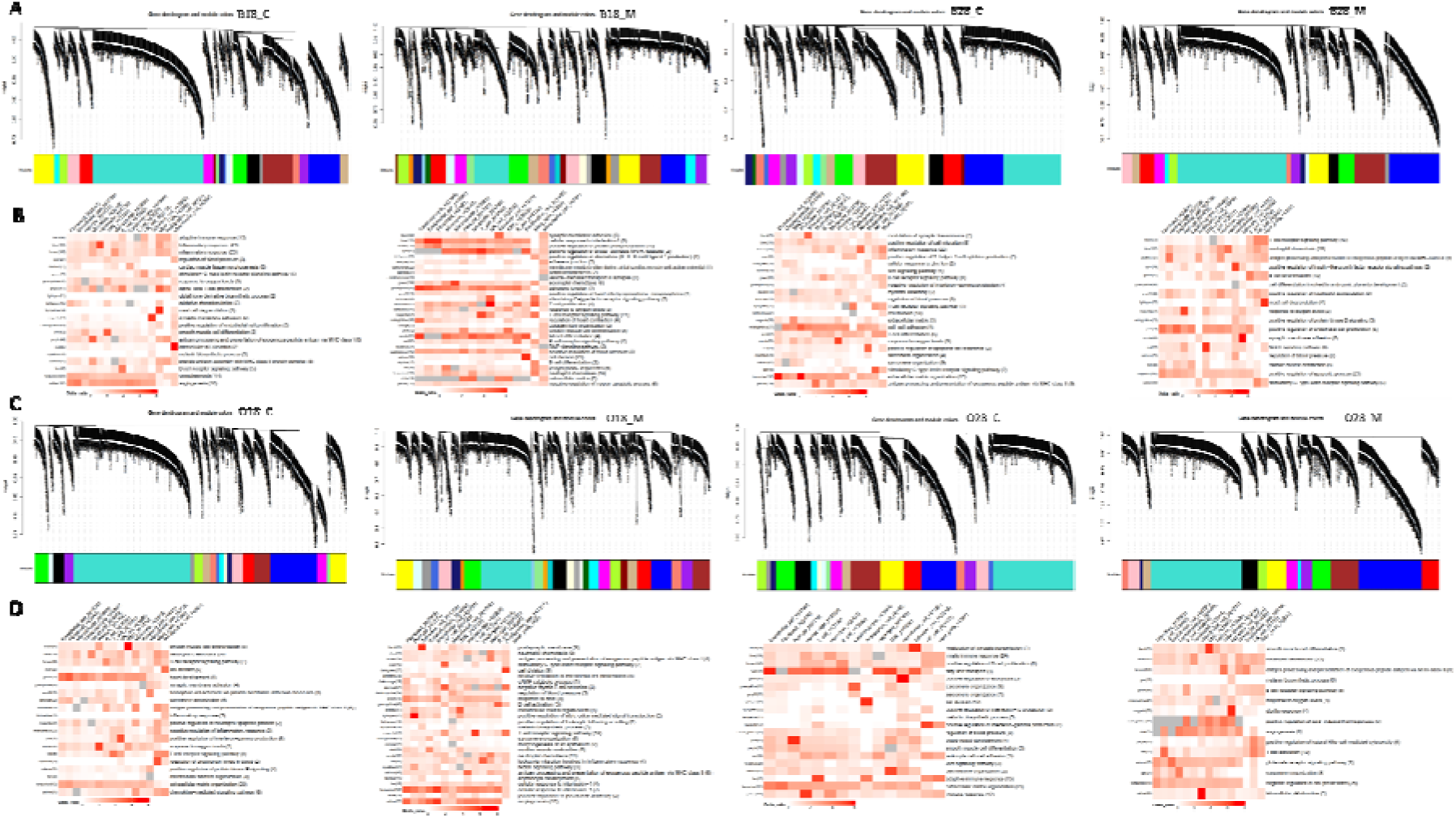
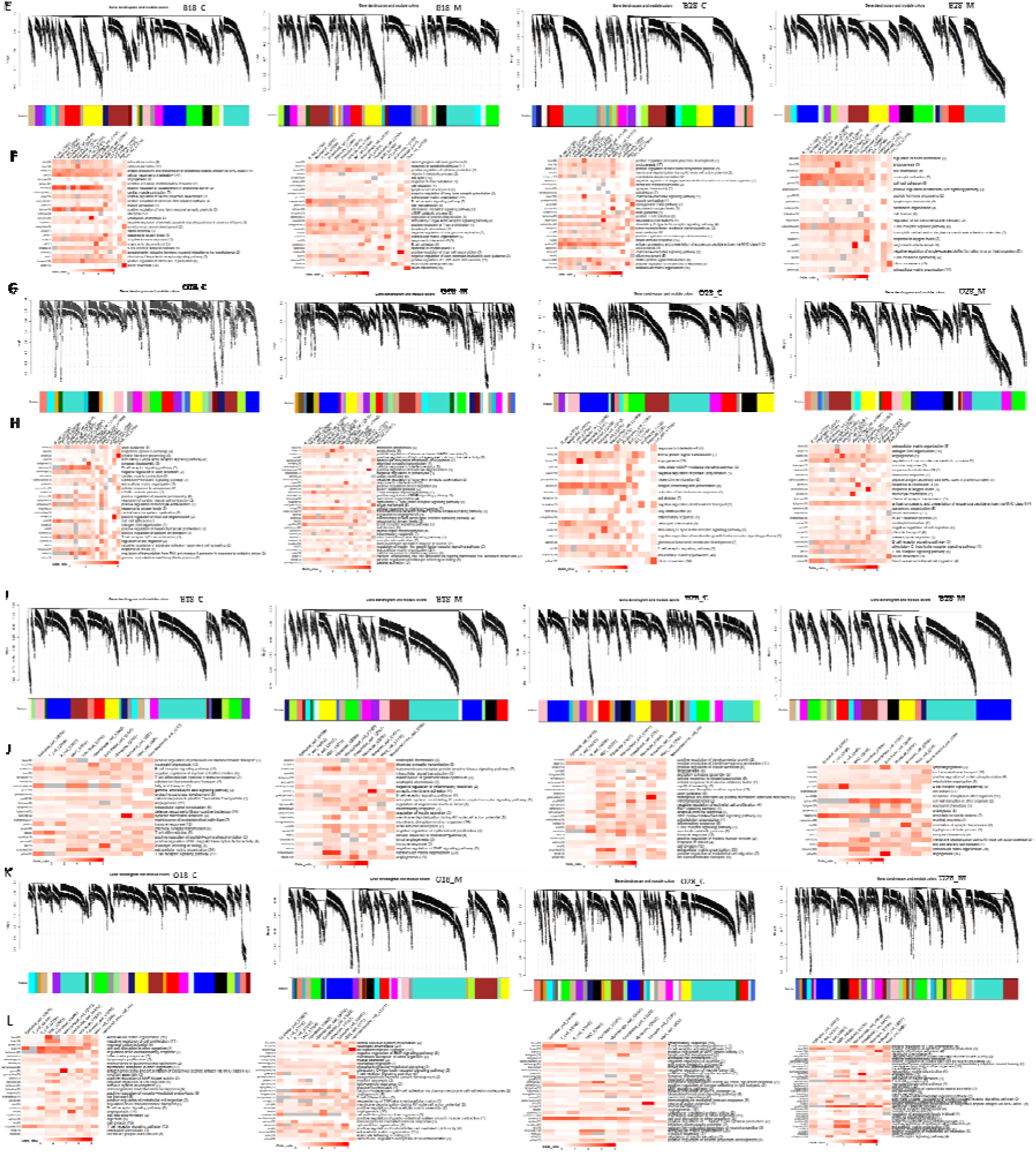
Identification and characterization of transcription modules using Weighted Gene Co-expression Network Analysis analysis in heart, lung and intestine in experimental animals. Cluster dendrograms of transcripts generated in the heart (A, C), lung (E, G) and intestine (I, K) as a result of WGCNA implementation in BSA and OVA model. Facet label above dendrograms represents BSA (B) or OVA (O), phase I (18) or phase II (28) and control (c) or model (M). Separation of transcription module is indicated by different colors. Gene enrichment analysis in each major cell type in heart (B, D), lung (F, H) and intestine (J, L) were analyzed by referring the public databases.

In the lung (Fig 7F), B cell receptor signaling was substantially higher in the model (OR = 5.90 in B cells) than in the control (OR = NA, data lacking), according to immunological and inflammatory responses. Stronger T cell activation was indicated by T cell receptor signaling, which displayed higher ORs in model T cells (OR = 5.23) compared to control (OR = 4.61). Compared to the control (OR = 1.20), lymphocyte chemotaxis was more noticeable in the model (OR = 3.82 in monocytes). Dynamic structural alterations were suggested by tissue remodeling and repair responses, which revealed that extracellular matrix organization was moved to SMCs in the model (OR = 2.74) but enriched in fibroblasts in the control (OR = 4.21). Vascular adaptation was demonstrated by the increased angiogenesis in the model’s endothelial cells (OR = 5.92) as compared to the control group (OR = 0.87). Cell-specific adaptations demonstrated that SMCs in the model responded more strongly to oxygen levels (OR = 5.78), as opposed to control (OR = 4.60), establishing a connection between hypoxia and the advancement of the disease.

Club cells in both groups had significantly higher levels of cilium movement, which is essential for lung function (model: 6.53; control: 5.94). However, in phase II model, immune activation showed that in contrast to control (OR = 3.70), neutrophil activation was significantly enriched in the model (OR = 5.21 in neutrophils), suggesting ongoing inflammation. The model’s macrophages presented more antigen via MHC class II (OR = 4.75) than the control group (OR = 3.50). Sarcomere organization was lacking in the control but increased in the model’s cardiomyocytes (OR = 4.45), according to structural and functional related study, indicating cardiac stress responses. Fibroblasts’ extracellular matrix structure was lower in the model (OR = NA) but remained high in the control group (OR = 4.12), suggesting phase-specific remodeling. Interleukin-6 was remarkably raised in AT2 cells in both the model (OR = 6.87) and control (OR = 6.66), indicating a consistent inflammatory marker, according to metabolic and signaling pathway-related responses. Compared to the control (OR = 2.15), T cell receptor signaling was more active in the model (OR = 3.32 in B cells). However, in OVA model (Fig 6H), significant odd ratios for a number of gene ontology keywords across cell types were seen in the control group during Phase I. For example, “respiratory gaseous exchange” was prevalent in Club_cell_L (2.49) and AT1_L (0.98), whereas “axon guidance” showed high odd ratios in Neutrophil_L (2.03), SMC_L (2.72), and AT1_L (1.99). The “B cell receptor signaling pathway” was very active in macrophage_cell_L (4.21), monocyte_L (4.54), and NK_cell_L (4.19). The model group, on the other hand, showed clear trends.

Macrophage_cell_L had a high odd ratio for “monocyte chemotaxis” (5.25), but ionocyte cell_L had a high odd ratio for “endocytosis” (7.11). T_cell_L had a much higher “T cell receptor signaling pathway” (5.74), whereas Mast cell_L had the highest “positive regulation of mast cell degranulation” (8.25). During Phase II, the control group showed high odd ratios for terms such as “neutrophil chemotaxis” in Neutrophil_L (4.87), “antigen processing and presentation” in NK_cell_L (5.47), and “response to interleukin-6” in AT2_L (3.71). “Extracellular matrix organization” was used frequently in Macrophage_cell_L (2.82) and Fibroblast_L (3.18). But for “response to interleukin-6” in AT2_L (5.72), “monocyte chemotaxis” in Macrophage_cell_L (6.08), and “response to oxygen levels” in SMC_L (6.48), the model group displayed higher odd ratios. In Neutrophil_L (3.24) and Proliferative_cell_L (3.12), the “B cell receptor signaling pathway” was enriched.

In the intestine (Fig 7J), while T cell differentiation is significantly lower (5.33 in control vs. model NA), indicating immune suppression in the model system, the model shows increased activity in negative regulation of inflammatory response (epithelial cells: 2.16 vs. control NA), transmembrane receptor protein tyrosine kinase signaling (SMCs: 3.69 vs. control 1.31), and synaptic membrane adhesion (Schwann cells: 6.38 vs. control 7.43) in Phase I. Phase II shows even more pronounced changes, with the model exhibiting significantly higher xenobiotic metabolic processes (epithelial cells: 5.68 vs. control NA) and cell wall disruption (SMCs: 5.05 vs. control 0), as well as robust modulation of synaptic transmission (Schwann cells: 5.68 vs. control 1.79). This reflects a shift toward metabolic adaptability and long-term neuromodulation during the chronic phase. B cell receptor signaling is differentially regulated (6.31 in model vs. 4.58 in control B cells), and extracellular matrix organization remains high but phase-specific (SMCs: 5.99 model vs. 5.40 control in Phase II). Most remarkably, inflammatory responses in macrophages peak differentially across phases (4.57 control vs. 3.47 model in Phase II), while angiogenesis in endothelial cells surges in the Phase II model (6.64) in contrast to control (NA). With continuously high synaptic activity in both phases (Phase I: 7.43 control vs. 6.38 model; Phase II: 5.68 model vs. lower control values), Schwann cells’ neuromodulatory function is especially clear. However, in the OVA (Fig 7L), Phase I of the model condition demonstrates a shift from generalized inflammation (control monocytes OR: 3.76 for inflammatory response) to more specific immune activation, as demonstrated by sustained B cell receptor signaling (OR increasing from 4.22 to 4.36) and increased T cell proliferation (macrophages OR: 3.75). From Phase I to Phase II (SMCs OR: 4.80 in control to 4.39 in model), structural remodeling processes, in particular extracellular matrix organization, intensify, indicating progressive tissue reorganization. Phase II exhibits clear biological changes, including a considerable increase in neutrophil chemotaxis (monocytes OR: 3.16, macrophages OR: 4.08), which suggests ongoing inflammatory activity, and angiogenesis in endothelial cells that is continually raised (OR: 5.02 in control to 5.71 in model). The model condition notably highlights nervous system processes, with Schwann cells exhibiting increased myelination (OR: 5.17) and neuroblast migration (OR: 1.31), and neuroendocrine cells becoming more active (OR: 2.76). The biological continuum from early immune responses to later tissue repair and neural adaption processes is reflected in these alterations.

## 3. Discussion

Food allergy (FA), a complex interplay between immune and effector cells initiates tightly coordinated allergic march that might result in life-threatening AS [16]. Traditionally, treatment solely relied on avoidance and emergency treatments due to accidental exposures. Currently, advancement in therapeutic research escalated the treatment options from avoidance to oral immunotherapy, sub-lingual immunotherapy, epicutaneous immunotherapy, modified allergens, DNA vaccines etc [17–21]. Apart from these, IgE monoclonal antibodies omalizumab and ligelizumab are in phase 3 clinical trials [22, 23]. However, a major area of unmet need stems from the fact that FA mediated pathological response is tailored to the allergenicity of each allergen, organ specific cellular microenvironment and sensitivity of an individual. Hence, understanding the intricate interactions between the cells in different organs during sensitization and shock phase will provide key insights to decipher complex multi-organ pathological response. Various multiomics and multi-tissue analysis in rats showed detailed interaction between immune cells, the change in mitochondrial and metabolic pathways across different organs and their importance in human health [24]. In recent years, various research groups use single-cell RNA sequencing (scRNA Seq) to understand the cell-specific molecular mechanisms underlying different allergies [25]. Leung et al., 2019 used skin transcriptome of atopic dermatitis (AD) with food allergy (FA) and reported that gene expression of dendritic cells and type 2 immune pathways are significantly increased compared to AD without FA. Single-cell RNA-sequencing of skin and peripheral blood mononuclear cells from AD patients showed differential immunological signatures that identify erythema and papulation, the two primary, qualitatively different skin manifestations of AD [26]. In a similar single-cell AD study, ovalbumin sensitization in a murine model caused Th2 cytokine genes expression to be upregulated in CD4+ T cells and mast/cell basophils, as well as an expansion of T cells, macrophages, dendritic cells (DCs), fibroblasts, and myocyte cell clusters in the skin. In this study, we evaluated the allergenicity at cellular level between two different allergens namely Bovine Serum Albumin (BSA), 69.2 kilodalton protein belonging to Serum Albumin family (from beef or cow milk) and Ovalbumin (OVA), a 42.8 kilodalton protein belonging to Serpin family (from Chicken egg white) [27]. Preliminary BLAST (Basic Local Alignment Search Tool) analysis showed no significant similarities in these two allergen protein sequences. Similar to previous reports, we reproduced FA model which was confirmed by significant increase in total IgE and allergen-specific IgE and occurrence of immediate anaphylaxis resulting to systematic death in our experimental animals [28]. The key broad findings from our study are (i) OVA causes stronger immune responses compared to BSA, (ii) intestine evokes a stronger immune response than heart and lungs, (iii) the signature of cellular activation is completely different in anaphylactic and sensitization phase (iv) cellular activation profile is organ-dependent even though the biological system is exposed to single allergen and (v) allergic phase mostly involves extensive stromal-immune cell interaction and shock phase involves myeloid-vascular cell cross-talk.

Detailed multi-organ cellular examination shows diverse but interrelated immune responses to food allergens, inferring a coordinated systemic reaction. In the heart, both BSA and OVA models demonstrated vascular dysfunction (pericyte loss: B28_M 749 cells, −29%; V28_M 503 cells, −47%) and cardiomyocyte adaptation (BSA: B28_M 2,199 cells, +147%), which is similar with IL-6-mediated cardiac remodeling observed in systemic inflammation [29, 30]. The lungs showed allergen-specific polarization, BSA drove B cell/T cell dominance (B28_M B cells: 13,109, +381%), whereas OVA prompted neutrophilia (V28_M: 10,275 cells, +326%), mimicking IL-17/IL-8 interaction reported in pulmonary allergy [31]. Surprisingly, OVA promoted fibroblast activation (V18_M: 4,259 cells, +840%) and Paneth cell expansion (+7,250%), implicating TGF-β-dependent fibrosis in the intestine, while BSA-induced CD8+ T cell infiltration (B18_M: 3,494 cells, +290%) correlated with decreased absorptive cells (−41%), similar to IFN-γ-driven barrier disruption [32, 33]. Intestinal inflammation may spread to distant organs by circulating cytokines (e.g., IL-6, IL-17), as shown in “leaky gut” disorders, according to the gut-cardio-pulmonary axis suggested by the shared neutrophilia (heart/lung) and IL-17 upregulation [34].

Furthermore, we found a common inflammatory core dominated by NOD-like receptor, cytokine-cytokine receptor interaction, TNF-α, NF-kB and JAK-STAT signaling all organs. These pathways are involved in innate immune sensing, amplification of cytokines, activation of endothelium, and recruitment of immune cells that are directly related to vascular leakage, hypotension and organ dysfunction in anaphylaxis [35–37]. Early stimulation at D18 and significant amplification at D28 propose that therapeutic targeting of these pathways during the allergic phase could prevent progression to systemic shock. These pathways (not specifically in FA/AS) have been reported as druggable targets such as TNF-α, JAK kinases and inflammasome associated pathways, suggesting avenues for preventative or adjuvant treatment. In the lung, transcriptional remodeling was dominated by chemokine signaling, Th17 differentiation and asthma associated pathways particularly during AS. This profile corresponds with the intense leukocyte recruitment and airway inflammation which is an important contributor to respiratory compromise in anaphylaxis. The enrichment of IL-17-related pathways identifies Th17 signaling as a potential amplifier of severe allergic inflammation over Th2 classical pathways [38–40]. We also observed significant increase in neutrophils in the model which might be due to Th17 inflammatory response [41, 42]. Targeting chemokine axes or IL-17-mediated responses may ameliorate pulmonary dysfunction and improve outcome during the systemic allergic reactions. Cardiac tissue was found to be accompanied by progressive inflammatory activation associated with the enrichment of metabolic and stress response pathways. These signatures are probably representative of hypoxia, oxidative stress and immune-mediated myocardial dysfunction during systemic anaphylaxis [43–45]. The intestine became a significant therapeutic nexus and the first and most varied immune activation. Enrichment of Th1/Th2 differentiation, IgA production, leukocyte transendothelial migration and tight junction pathway suggest the concurrent activation of adaptive immunity, effector responses IgG-mediated and epithelial barrier disruption. The marked increase of inflammatory pathways by D28, especially the OVA model, agrees with a model of loss of mucosal immune regulation and barrier integrity as a prerequisite for systemic spread of the allergen and immunological amplification of the immune response [46–48]. These results indicate that interventions towards restoring epithelial barrier integrity or modulating immune responses located in the gut may provide a significant impact in preventing severe systemic anaphylaxis.

Therapeutic strategies focused on FcεRI signaling, epithelial tight junctions or mucosal cytokine networks may therefore be efficient points of early intervention. The involvement of inflammatory cells as well as metabolic pathway would indicate that cardioprotective strategies that use a combination of immune modulation and metabolic support may be required to prevent or limit cardiac injury from severe anaphylaxis. Few therapeutic strategies such as the use of mesenchymal stem cells, nutraceuticals have been reported to modulate immune response as well as metabolic support [49–51]. In addition to shared inflammatory mechanisms, OVA induced larger and sustained pathway activation than BSA across all organs at D28, indicating allergen-dependent differences in immunogenicity and systemic impact. These results suggests that therapeutic intervention timing should be tailored to allergen-specific risk profiles, an important consideration for personalized management of food allergy. Pathway trajectory analysis further revealed a temporal shift in which early inflammatory signaling precedes widespread immune and metabolic dysregulation. This progression provides a potentially important time for intervention before the development of irreversible systemic pathology, the so-called anaphylactic phase. In conclusion, our multi-organ pathway analysis revealed conserved inflammatory hubs and tissue-specific immune programs that collectively drive food allergy–induced anaphylaxis. These data support a therapeutic model in which early targeting of common innate and cytokine-driven pathways; in particular at the gut-immune interface, may prevent progression to systemic, life-threatening disease. Such strategies may provide a complement to current emergency treatments and take the management of anaphylaxis out of the realm of emergency treatment and into prevention and risk reduction.

As mentioned earlier, AS is a multi-cellular response, which is synchronized through ligand-receptor interactions between immune cells, endothelial cells, and tissue-resident cells. Therefore, to analyze these interactions, the cell-chat approach allowed us to relate the intracellular alterations of the pathway found by the KEGG analysis to the extracellular signal transduction found by CellChat, and this approach allowed us to provide an overall framework of how cell-intrinsic transcriptional changes are related to intercellular crosstalk in the presence of allergen-induced organ dysfunction. Mirroring the shift from localized inflammation to systemic responses, Phase I of the BSA model exhibits dominating fibroblast-macrophage interactions (weight=3.27), whereas Phase II transitions to robust fibroblast-neutrophil crosstalk (weight=2.69) [16]. The OVA model has strong fibroblast-Schwann cell connections in Phase I (O18M, weight=3.06), indicating neuroimmune involvement in early allergy symptoms [52]. In Phase II, fibroblast-neutrophil communication is high (O28M, weight=1.79). Serosal surfaces are implicated in allergic inflammation via both allergens’ persistent macrophage-mesothelial cell contacts (BSA: weight=1.75; OVA: weight=1.32) [53]. Significantly, fibroblast-endothelial cell interactions are higher in the OVA model (weight=1.83 compared to BSA’s 1.27), which may account for variations in the degree of vascular leakage caused by different allergens [54]. These results point to allergen-specific differences in cell communication networks and conserved pathways (fibroblast-immune-neuro axis) that may help guide targeted treatments for various food allergies.

In lungs, Phase I (food allergy) and Phase II (anaphylaxis) reactions exhibit different patterns when cell-cell interactions in lung tissues are analyzed. We found fibroblast-immune cell interactions predominate in Phase I, especially between Fibroblast and Macrophage cell (weight: 3.97 in BSA 18M), which reflects tissue remodeling and chronic inflammation typical of allergic sensitization [31]. Moderate interactions between adaptive immune cells, such as T cell and B cell, are consistent with early immunological activation and allergen identification. The histamine-mediated vascular permeability and bronchoconstriction of anaphylaxis are mirrored in Phase II, which shows a dramatic shift toward vascular and effector cell responses. Mast cells show noticeably stronger interactions with Endothelial cells (weight: 0.52 in BSA 28M) and SMC [35]. Phase II also shows an increase in neutrophil contacts, indicating that the creation of neutrophil extracellular traps plays a role in systemic inflammation [55]. In Phase II, mast cells/endothelial-mediated systemic responses replace fibroblast-driven localized inflammation, offering new mechanistic insights into the development of allergies. The results point mast cell stabilization as a potential target for reducing the severity of anaphylaxis and fibroblast-immune crosstalk as a potential target for preventing the development of allergies, providing new treatment approaches for these clinically different stages of allergic illness.

Analyzing intestinal cell-cell interactions in food allergy (Phase I) and AS (Phase II) revealed different patterns. Strong interactions between immunological cells (such as T cell, B cell, and Macrophage) and structural cells (such as Epithelial and Fibroblast) were noted in Phase I (e.g., BSA 18M and OVA18M). Strong immune-stromal crosstalk was demonstrated in BSA 18M, where SMC-macrophage cell had the highest interaction weight (2.627) and fibroblast-macrophage (1.727). Similarly, SMC-macrophage displayed the highest weight (2.762) in OVA 18M, indicating inflammation mediated by macrophages. These results are consistent with research showing how immune activation impairs barrier function in food allergy development through interactions between macrophages and epithelium [56]. Systemic participation was indicated by interactions shifting toward endothelial and Schwann cells in Phase II (e.g., B28M and O28M).

Schwann cel-SMC, for instance, exhibited the highest weight (5.295) in B28M, suggesting that neuro-immune crosstalk plays a role in anaphylaxis. This is consistent with reports of severe allergic reactions involving neural regulation [57]. OVA 28M displayed increased SMC-schwann connections (2.438), indicating that neuromuscular signaling may play a new function in anaphylactic shock. This study highlights understudied pathways such as Schwann cell participation and depicts dynamic transitions in cell-cell communication from localized allergy (Phase I) to systemic anaphylaxis (Phase II) through unexplored mechanisms. Targeting neuro-immune late pathways and macrophage-epithelial early interactions may slow the progression of the disease, according to the study.

This multi-organ cell chat analysis established particular ligand/receptor combinations and signaling pathways that mediate intercellular communication between different cell types in the process of allergic sensitization and development of anaphylactic shock. In order to further explain these transcriptional programs that we observed between these cell-cell interactions, we conducted WGCNA to define the modules of co-expressing genes and hub genes that orchestrate cellular responses to paracrine signaling at different phases of disease and in different tissue conditions. This combination gave us the possibility to connect the extracellular communication architecture to intracellular gene control systems and understand how intercellular communication leads to coordinated transcriptional response in multi-organ dysfunction. According to the results of our WGCNA study of the heart, lung, and intestine in BSA and OVA food allergy mice models, Phase I (allergic phase) and Phase II (anaphylactic shock phase) show dynamic immunological, structural, and metabolic changes. The model demonstrated a strong surge of immune activation at shock phase (day 28) characterized by a significant upregulation of the neutrophil recruitment (OR = 2.87), and dietary immune activation: T cell receptor signaling (OR = 5.28). The increased response of T cells, macrophags, and neutrophils indicates an increase in the inflammatory state, which is a key characteristic of anaphylaxis and allergic responses. Interestingly, macrophages and neutrophils were found more engaged in antigen processing and presentation (OR = 1.30 vs. 4.75 in the model), which points to the activation of the immune system that is antigen recognition and response oriented at the shock stage. Activation of mast cells (OR = 6.21) also implied that degranulation and the release of histamine are vital elements in the further development of the inflammatory response [58]. Vascular remodeling, especially in the endothelial cells was seen in several organs. Endothelial cells and fibroblasts were found to be more enriched in the heart with “positive regulation of endothelial cell proliferation” (OR = 5.00) indicating that the vascular repair and remodeling processes are underway in the heart in response to the injury. This augmented neutrophil recruiting may also indicate a new understanding of the role of neutrophil in the augmentation of inflammation in the heart amid shock and might be exploited to treat this condition. Immunological cell recruitment remained a vital part of the immune response in the lung, although in the model, the T cell receptor signaling was more active (OR = 5.23). Also, the enhanced activities of the neutrophil chemotaxis (OR = 3.50) and macrophage antigen presentation (OR = 4.75) serve to highlight the prolonged immune stimulation that occurs in the lung at the shock stage. Those immunological reactions can cause a worsening of tissue damage and lead to airway narrowing and inflammation, which are characteristic of AS and asthma-like conditions triggered by allergies [59]. The augmented angiogenesis in the lung in the model (OR = 5.92 in endothelial cells) could be an adaptive reaction in the lung to support the provision of oxygen to the damaged tissue, which aligns with the concept of the compensatory vascular alterations to the chronic inflammation and the hypoxia [60]. Another significant innovativeness in the lung was the high rate of increase in the endothelial cells angiogenesis (OR = 5.92) was found to be much higher in the model than the control group. This means that in reaction to AS, lung endothelial cells can undergo adaptive angiogenesis to enhance the perfusion of the tissues and supply of oxygen. It is a new understanding of how lungs react to allergic responses, namely, this adaptive mechanism of coping with vascular stress (Wang et al., 2021). Moreover, macrophages activation in the lung through MHC class II is higher (OR = 4.75) and this highlights the role of the lung in antigen processing in the shock phase, which can lead to the protracted inflammation and tissue injury. An interesting change was observed in the intestine where instead of the model exhibiting immune suppression (reduced T cell differentiation) there was activation of metabolic processes and this included the xenobiotic metabolic processes (OR = 5.68). This metabolic plasticity may be a response by the body to regain homeostasis perhaps by the elimination of inflammatory mediators and allergens. The enhancement of synaptic membrane adhesion and neuromodulation in Schwann cells (OR = 6.38) indicate that there is a neurological response to long-term stress, which may indicate the linkage of the immune system and neural networks to long-term inflammation [57]. The intestinal fibroblasts also experienced greater “extracellular matrix organization” (OR = 5.99), which is evidence of intestinal tissue repair mechanism that attempts to restore its barrier functions and prevent additional damage. These results show that tissue remodeling and vascular adaptation are important part of the recovery phase that indicates that therapies which focus on vascular repair and extracellular matrix reorganization may be useful in preventing tissue damage in the long term after AS. The results of the current research highlight a number of possible treatment plans to use in the treatment of food allergies and AS. The T cell and macrophage signaling pathway (e.g. “ T cell receptor signaling “) upregulation in different organs indicates that immune cells, especially T cells and macrophages, might be targeted to reduce excessive inflammation. The activation of neuromodulatory processes, in particular, Schwann cells, which had a significant increase in the synaptic membrane adhesion (OR = 6.38) and neuromodulation (OR = 5.68), was a novel finding in the intestine. It implies that in the case of chronic inflammation in anaphylaxis caused by food allergies, neural pathways can be engaged in the regulation of immune responses. Another new feature is the change in the immune suppression to metabolic adaptation, an increased metabolism of xenobiotics (OR = 5.68), which suggests that the intestine is trying to eliminate toxic agents or allergens in the event of chronic inflammation, which may become a possible target of therapeutic intervention. Based on our results therapeutic strategies focusing on the following might be crucial in preventing food allergy mediated anaphylactic shock-1) Immunomodulatory immunotherapies, including corticosteroids, monoclonal antibodies that block certain immune checkpoints (e.g., anti-IL-4, anti-IL-13), or T cell receptor-inhibitors, might be explored to suppress the immune response in anaphylaxis. 2) It is the fact that the degranulation of the mast cells is increased in the heart and the lung, which points to the importance of mast cells, which are essential to the allergic processes. Cromolyn sodium or antihistamines are used, but not efficient to prevent the acute phase of anaphylaxis due to the degranualtion of mast cells. 3) It is also an indication that through intensified angiogenesis of endothelial cells, the repair processes of the vessels might be useful in the recovery of tissues during chronic stage. Endothelial cell proliferation agents or vascular endothelial growth factor (VEGF) inhibitors could be helpful in regulating tissue injury and healing and 4) The alterations in Schwann cell activity and synaptic activity in the intestine indicate the involvement of neural pathways in immunity responses. Neuroimmune interactions and possibilities of neuroprotective strategies can be explored to present new therapeutic opportunities in the management of chronic inflammation and remodeling in food allergies. In summary, this WGCNA analysis indicates that the immune, vascular, and structural remodeling processes that take place during AS caused by food allergies are rather complex. With an emphasis on immune modulation, mast cell stabilization, vascular healing, and neuroimmune communications, future treatment approaches can be designed that can be more effective in the treatment of both acute anaphylaxis and chronic allergy.

## 4. Methods

### 4.1. Animal Sensitization

Male c57BL6 (SPF grade) mice were procured from Zhuhai Baishitong Biotechnology company and acclimatized for 1 week at SPF Laboratory Animal Center, Shenzhen LingFu TopBiotech. Co., LTD before starting the experiments. Animal experimental procedures were strictly carried out according to the guidelines approved by Topbiotech Biotechnology Co.,Ltd. SYXK (LFTOP-IACUC-2023-0013), Guangdong, China. Animals were randomly divided into 8 groups and had ad libitum access to food and water. The groups were as follows.

Group A: BSA control group (Pertussis toxin, 300 ng/100 gm) sacrificed at Phase I (day 18, n=4),

Group B: BSA (1mg) + Pertussis toxin (300 ng/100 gm) sensitized group sacrificied at phase I (day 18, n=4),

Group C: OVA control group (Aluminium hydroxide, 3.5 mg) sacrificed at phase I (day 18, n=4),

Group D: OVA (1 mg) + Aluminium hydroxide (3.5 mg) sensitized group sacrificed at phase I (day 18, n=4),

Group E: BSA control group (Pertussis toxin,300 ng/100 gm) sacrificed at Phase II (day 28, n=4),

Group F: OVA control group (Aluminium hydroxide, 3.5 mg) sacrificed at phase II (day 28, n=4),

Group G: BSA (1mg) + Pertussis toxin (300 ng/100 gm) sensitized group sacrificed at phase II (day 28, n=4),

Group H: OVA (1 mg) + Aluminium hydroxide (3.5 mg) sensitized group sacrificed at phase II (day 28, n=4).

The route of administration of adjuvants/allergens and the timeline for different groups are described in Fig 1. According to the experimental timeline, animals were sacrificied and organs were collected at Phase I (Blood, Heart, Lung and Intestine) and Phase II (Blood, Heart, Lung and Intestine).

### 4.2. Enzyme-linked immunosorbent assay

Blood was collected from the experimental animals, plasma was separated and total-IgE (MM-0063R2 - Jiangsu Meimian Industrial Co., Ltd, Jiangsu, China), BSA-specific IgE (EM2289 - FineTest) and OVA-specific IgE (MM-20974R1 - Jiangsu Meimian Industrial Co., Ltd, Jiangsu, China) were quantified according to the manufacturer’s instructions.

### 4.3. Tissue dissociation and cell purification

Tissues (PBMC, heart, lung and intestine) were transported in sterile culture dish with 10 ml 1x Dulbecco’s Phosphate-Buffered Saline (DPBS; Thermo Fisher, Cat. no. 14190144) on ice to remove the residual tissue storage solution, then minced on ice. We used dissociation enzyme 0.25% Trypsin (Thermo Fisher, Cat. no.25200-072) and 10 ug/mL lDNase I (Sigma, Cat. no. 11284932001) dissolved in PBS with 5% Fetal Bovine Serum (FBS; Thermo Fisher, Cat. no. SV30087.02) to digest the tissues. Lung, heart, intestine and peripheral blood mononuclear cells were dissociated at 37 with a shaking speed of 50 r.p.m for about 40 min. We repeatedly collected the dissociated cells at interval of 20 min to increase cell yield and viability. Cell suspensions were filtered using a 40 um nylon cell strainer and red blood cells were removed by 1X Red Blood Cell Lysis Solution (Thermo Fisher, Cat. no. 00-4333-57). Dissociated cells were washed with 1x DPBS containing 2% FBS. Cells were stained with 0.4% Trypan blue (Thermo Fisher, Cat. no. 14190144) to check the viability on Countess® II Automated Cell Counter (Thermo Fisher). The sample was then sent to Majorbio Bio-pharm Technology Co., Ltd (Shanghai) for testing.

### 4.4. 10x library preparation and sequencing

Beads with unique molecular identifier (UMI) and cell barcodes were loaded close to saturation, so that each cell was paired with a bead in a Gel Beads-in-emulsion (GEM). After exposure to cell lysis buffer, polyadenylated RNA molecules hybridized to the beads. Beads were retrieved into a single tube for reverse transcription. On cDNA synthesis, each cDNA molecule was tagged on the 5’end (that is, the 3’end of a messenger RNA transcript) with UMI and cell label indicating its cell of origin. Briefly, 10× beads that were then subject to second-strand cDNA synthesis, adaptor ligation, and universal amplification. Sequencing libraries were prepared using randomly interrupted whole-transcriptome amplification products to enrich the 3’ end of the transcripts linked with the cell barcode and UMI. All the remaining procedures including the library construction were performed according to the standard manufacturer’s protocol (Chromium Single Cell 3 v3.1). Sequencing libraries were quantified using a High Sensitivity DNA Chip (Agilent) on a Bioanalyzer 2100 and the Qubit High Sensitivity DNA Assay (Thermo Fisher Scientific). The sequencing were sequenced on Novaseq Xplus using PE150 mode.(Novaseq Xplus DNBSEQ-T7). The sequencing libraries were performed on DNBSEQ-T7 platform (PE150) using DNBSEQ-T7RS Reagent Kit (FCL PE150) version 3.0. The sequencing and bioinformatic analysis were performed on platform of Majorbio Co., Ltd (Shanghai, China).

### 4.5. Single cell RNA-seq data processing

Reads were processed using the Cell Ranger (v7.1.0) pipeline with default and recommended parameters. FASTQs generated from Illumina sequencing output were aligned to the mouse genome, version GRCm38, using the STAR algorithm [61]. Next, Gene-Barcode matrices were generated for each individual sample by counting UMIs and filtering non-cell associated barcodes. Finally, we generate a gene-barcode matrix containing the barcoded cells and gene expression counts. This output was then imported into the Seurat (v5.0) R toolkit for quality control and downstream analysis of our single cell RNAseq data[62]. All functions were run with default parameters, unless specified otherwise. We first filtered the matrices to exclude low-quality cells using a standard panel of three quality criteria: (1) number of detected transcripts (number of unique molecular identifiers); (2) detected genes; and (3) percent of reads mapping to mitochondrial genes (The numbers out of the limit of mean value +/- 2-fold of standard deviations). The expression of genes was calculated using Percentage Feature Set function of the seurat package [62]. The normalized data (NormalizeData function in Seuratpackage) was performed for extracting a subset of variable genes. Variable genes were identified while controlling for the strong relationship between variability and average expression. Next, we integrated data from different samples after identifying ‘anchors’ between datasets using FindIntegrationAnchors and IntegrateData in the seurat package [62, 63]. Then we performed principal component analysis (PCA) and reduced the data to the top 30 PCA components after scaled the data. We visualized the clusters on a 2D map produced with t-distributed stochastic neighbor embedding (t-SNE) [64].

### 4.6. Identification of cell types and subtypes by nonlinear dimensional reduction (t-SNE)

Cells were clustered using graph-based clustering of the PCA reduced data with the Louvain Method [65] after computing a shared nearest neighbor graph [62]. For sub-clustering, we applied the same procedure of scaled, dimensionality reduction, and clustering to the specific set of data (usually restricted to one type of cell.) For each cluster, we used the Wilcoxon Rank-Sum Test to find significant deferentially expressed genes comparing the remaining clusters. SingleR [66] and known marker genes were used to identify cell type.

### 4.7. Differential expression analysis and Functional enrichment

To identify DEGs (differential expression genes) between two different samples or clusters, analysis was performed using the function FindMarkers in Seurat, using a likelihood ratio test. Essentially, DEGs with |log2FC| > 0.25 and Q value ≤ 0.05 were considered to be significantly different expressed genes. In addition, functional-enrichment analysis GO were performed to identify which DEGs were significantly enriched in GO terms and metabolic pathways at Bonferroni-corrected P-value ≤ 0.05 compared with the whole-transcriptome background. GO functional enrichment analysis were carried out by Goatools (https://github.com/tanghaibao/Goatools).

### 4.8. Cytokine-receptor coupling

CellChat was used to predict cell-cell interactions by analyzing the expression of known ligand-receptor pairings in various cell types. To identify potential cell-cell communication networks altered or provoked in our experimental samples, we followed the recommended workflow and loaded the normalized counts into CellChat, where we used the preprocessing functions identifyOver Expressed Genes and identify over expressed Interactions between the ligand and receptors of various cells with standard set parameters. Furthermore, to identify the senders and receivers in the network, the function named netAnalysis_signalingRole was used on the netP data slot [67].

### 4.9. Weighted Gene Co-expression network analysis

Weighted Gene Co-expression network analysis was performed to understand the transcriptome-wide relationship based upon pairwise correlation between variables (modules) as per previous publication [68]. The R package ‘hdWGCNA’ uses weighted gene co-expression network analysis (hdWGCNA) to create a scale-free network at the single cell level. To achieve optimal connectivity, the scale-free topology model fit threshold was set at >0.85, and a soft threshold of 5 was selected. GSVA was utilized to evaluate the TCGA cohort with modules. The Spearman test was used to analyze correlations between modules and phenotypes.

## Reporting Summary

The current paper is the first single-cell resolution map of the pathogenesis of food allergy in three vital organs in both sensitization and AS phase. We have discovered basic understanding of how various allergens trigger unique pathogenic pathways through the application of integrated single-cell transcriptomics, intercellular communication analysis and network biology methods and found that each allergen presents unique and organ-specific patterns of vulnerability to distinct pathogeneses.

One observation derived in our study was that OVA consistently produced higher immune responses than BSA and the intestine was the most dramatically remodeled cellular tissue, the lung was polarized to antigenic immune responses, and the heart showed severe vascular injury with a remarkable adaptive capacity maintained by adaptive mechanisms. These organ-specific reactions underscore the nature of systemic allergic reactions and complicate the idea of standardized treatment of allergies.

The phase changes were marked by dynamic reprogramming of the molecules, and the development of the inflammasome signal and the fracture of the anti-inflammatory processes of their own origin in anaphylaxis. The conserved inflammatory mechanisms between the organs, such as a NOD-like receptor, cytokine-cytokine receptor, TNF-α, and NF-kappa B pathways, indicates that this form the events that can be targeted by drugs to prevent the transition of the sensitizing state into the systemic shock.

This research identified three key conceptual steps. One, it provides the first multi-organ cellular roadmap of FA progression displaying organ-specific, but coordinated responses that require multi-targeted therapeutic responses. Second, it establishes the involvement of previously unknown cells such as the role of erythrocytes in allergic inflammation and reveals the essential neuroimmune mechanisms of Schwann cells that potentially represent new intervention targets to prevent the escalation of anaphylactic responses. Third, it shows allergen-specific patterns of endothelial failure and dysfunction of the barrier, indicating that there is a need to treat individuals in a personalized approach, depending on specific allergen profiles.

The cell-cell interaction networks that we have determined show a profound temporal transition of localized fibroblast-immune crosstalk in sensitization to systemic neuro-vascular interactions in anaphylaxis. The evolution of Schwann cell-mediated signalling in severe reactions creates completely novel pathways of neuromodulatory intervention that were previously not a part of anaphylaxis management.

The network analysis of the evolving patterns of early-stage extracellular matrix remodeling to late-stage synaptic modulation and metabolic changes has shown us a time-scale of the disease development. This implies that the timing of therapeutic intervention should be specific to each stage of the disease process, where early interventions are used to target tissue remodelling and barrier integrity whilst late-stage interventions may be useful in counteracting neuro-vascular dysfunction and metabolic stress.

There are a number of translational opportunities arising out of this work. Disease progression and treatment response biomarkers can be developed because of the identification of phase-specific molecular signatures. The patterns of intestinal immune activation indicate that gut barrier composition and mucosal immune control methods can play a disproportionate contribution to the prevention of the systemic dissemination of allergens. The cardiac compensatory responses we observed imply that aids to such compensatory responses may prevent cardiovascular failure in acute responses.

The allergen-specific pathways that we have identified should be pursued in the wider allergen panels and patient cohorts to determine their clinical utility. The future research perspectives involve validation of the studies in human tissue and functional research studies to evaluate the therapeutic utility of the target of neuro-vascular linkage, the investigation of the long-term structural after-effects in the organs involved, and the development of the multi-organ protection strategies considering coordination of the systemic allergic responses.

In conclusion, the power of this single-cell atlas is the primary contribution to the knowledge of food allergy pathogenesis since it demonstrates the complexity of intercellular and intracellular interactions of immune cells, structural cells, and neural components in various organs. It offers a platform on the development of next-generation treatment plans that go beyond emergency responses to prevention with multi-organ protection, phase-specific response, and neuroimmune modulation to avoid life-threatening anaphylaxis.

## 6. Data availability & code availability statement

Inquiries about data can be directed to the corresponding author.

## Acknowledgements

This study was funded by the following grant: National Key R&D Program Intergovernmental Key Project: 2023YFE0114300.

## 7. Authors Contributions

R.L.J drafted the manuscript, analyzed the data and generated the figures. X.X and M.A.A analyzed the data and interpreted the figures. Y.L, S.M, S.K, and A.N performed the experiments. A.B, did the conceptualization of the research, designed and supervised the experiment, critical revision, obtained the fundings. X.L, participated in fundings and critically reviewed the manuscript. J.X, C.L, assisted in procurement animal maintenance and preparation for the experiments. S.L.P, B.H, and M.A contributed ideas and expertise. All authors revised and edited the manuscript.

## 8. Competing interests

No competiting interests

## 9. Additional information

Nil

## Notes

### Competing Interest Statement

The authors have declared no competing interest.

## References

1. Ozdemir, C., et al., Lifestyle Changes and Industrialization in the Development of Allergic Diseases. Current Allergy and Asthma Reports, 2024: p. 1–15.

2. van Ree, R., et al., The COMPARE Database: A Public Resource for Allergen Identification, Adapted for Continuous Improvement. Frontiers in Allergy, 2021. Volume 2 - 2021.

3. Pomés, A., Intrinsic properties of allergens and environmental exposure as determinants of allergenicity. Allergy, 2002. 57(8): p. 673–679.

4. Dearman, R.J., et al., Divergent antibody isotype responses induced in mice by systemic exposure to proteins: a comparison of ovalbumin with bovine serum albumin. Food Chem Toxicol, 2000. 38(4): p. 351–60.

5. Kanagaratham, C., et al., IgE and IgG Antibodies as Regulators of Mast Cell and Basophil Functions in Food Allergy. Front Immunol, 2020. 11: p. 603050.

6. Gbenga-Olusanya, O., et al., Mechanisms and Environmental Determinants of Food Allergies: An Integrative Review. Faculty of Natural and Applied Sciences Journal of Applied Biological Sciences, 2025. 2(3): p. 61–76.

7. Pucci, S. and C. Incorvaia, Allergy as an organ and a systemic disease. Clinical & Experimental Immunology, 2008. 153: p. 1–2.

8. Pucci, S. and C. Incorvaia, Allergy as an organ and a systemic disease. Clin Exp Immunol, 2008. 153 Suppl 1(Suppl 1): p. 1–2.

9. Devonshire, A., et al., Multi-omics profiling approach in food allergy. World Allergy Organization Journal, 2023. 16(5): p. 100777.

10. Turner, P.J., et al., Time to revisit the definition and clinical criteria for anaphylaxis? World Allergy Organ J, 2019. 12(10): p. 100066.

11. Al-Salam, S., et al., Cellular and immunohistochemical changes in anaphylactic shock induced in the ovalbumin-sensitized Wistar rat model. Biomolecules, 2019. 9(3): p. 101.

12. Bellou, A., et al., Combined Treatment with KV Channel Inhibitor 4-Aminopyridine and either γ-Cystathionine Lyase Inhibitor β-Cyanoalanine or Epinephrine Restores Blood Pressure, and Improves Survival in the Wistar Rat Model of Anaphylactic Shock. Biology, 2022. 11(10): p. 1455.

13. Bellou, A., et al., 4-Aminopyridine, a blocker of Voltage-dependent K+ channels, restores blood pressure and improves survival in the Wistar rat model of anaphylactic shock. Critical care medicine, 2016. 44(11): p. e1082–e1089.

14. Muraro, A., et al., Precision medicine in allergic disease-food allergy, drug allergy, and anaphylaxis-PRACTALL document of the European Academy of Allergy and Clinical Immunology and the American Academy of Allergy, Asthma and Immunology. Allergy, 2017. 72(7): p. 1006–1021.

15. Korsunsky, I., et al., Fast, sensitive and accurate integration of single-cell data with Harmony. Nat Methods, 2019. 16(12): p. 1289–1296.

16. Reber, L.L., J.D. Hernandez, and S.J. Galli, The pathophysiology of anaphylaxis. Journal of Allergy and Clinical Immunology, 2017. 140(2): p. 335–348.

17. Andorf, S., et al., Anti-IgE treatment with oral immunotherapy in multifood allergic participants: a double-blind, randomised, controlled trial. Lancet Gastroenterol Hepatol, 2018. 3(2): p. 85–94.

18. Andorf, S., et al., A Phase 2 Randomized Controlled Multisite Study Using Omalizumab-facilitated Rapid Desensitization to Test Continued vs Discontinued Dosing in Multifood Allergic Individuals. EClinicalMedicine, 2019. 7: p. 27–38.

19. MacGinnitie, A.J., et al., Omalizumab facilitates rapid oral desensitization for peanut allergy. J Allergy Clin Immunol, 2017. 139(3): p. 873–881.e8.

20. Sindher, S.B., et al., Phase 2, randomized multi oral immunotherapy with omalizumab ‘real life’ study. Allergy, 2022. 77(6): p. 1873–1884.

21. Takahashi, M., et al., Oral immunotherapy combined with omalizumab for high-risk cow’s milk allergy: a randomized controlled trial. Sci Rep, 2017. 7(1): p. 17453.

22. Fiocchi, A., B.P. Vickery, and R.A. Wood, The use of biologics in food allergy. Clin Exp Allergy, 2021. 51(8): p. 1006–1018.

23. Dantzer, J.A. and R.A. Wood, Omalizumab as an adjuvant in food allergen immunotherapy. Curr Opin Allergy Clin Immunol, 2021. 21(3): p. 278–285.

24. authors, M.S.G.P., et al., Temporal dynamics of the multi-omic response to endurance exercise training. Nature, 2024. 629(8010): p. 174–183.

25. Mitamura, Y. and C.A. Akdis, Single-cell analysis of allergic diseases. Allergy, 2023. 78(2).

26. Sekita, A., et al., Multifaceted analysis of cross-tissue transcriptomes reveals phenotype–endotype associations in atopic dermatitis. Nature communications, 2023. 14(1): p. 6133.

27. Radauer, C., et al., Allergens are distributed into few protein families and possess a restricted number of biochemical functions. J Allergy Clin Immunol, 2008. 121(4): p. 847–52.e7.

28. Schülke, S. and M. Albrecht, Mouse Models for Food Allergies: Where Do We Stand? Cells, 2019. 8(6).

29. Pinto, A.R., et al., Revisiting Cardiac Cellular Composition. Circ Res, 2016. 118(3): p. 400–9.

30. Preda, A., et al., IL-6 in the spotlight: From cardiovascular pathophysiology to therapy. Eur J Clin Invest, 2025: p. e70161.

31. Lambrecht, B.N. and H. Hammad, The immunology of asthma. Nat Immunol, 2015. 16(1): p. 45–56.

32. Peterson, L.W. and D. Artis, Intestinal epithelial cells: regulators of barrier function and immune homeostasis. Nat Rev Immunol, 2014. 14(3): p. 141–53.

33. Quintero, M., et al., Intestinal crypts remodel in response to deletion of Notch pathway ligands from Paneth cells. Physiology, 2025. 40(S1): p. 1026.

34. Zheng, P., et al., The gut microbiome modulates gut–brain axis glycerophospholipid metabolism in a region-specific manner in a nonhuman primate model of depression. Molecular psychiatry, 2021. 26(6): p. 2380–2392.

35. Galli, S.J., M. Tsai, and A.M. Piliponsky, The development of allergic inflammation. Nature, 2008. 454(7203): p. 445–454.

36. Pumphrey, R., Anaphylaxis: can we tell who is at risk of a fatal reaction? Curr Opin Allergy Clin Immunol, 2004. 4(4): p. 285–90.

37. Stone, K.D., C. Prussin, and D.D. Metcalfe, IgE, mast cells, basophils, and eosinophils. Journal of Allergy and Clinical Immunology, 2010. 125(2): p. S73–S80.

38. Lloyd, C.M. and S. Saglani, T cells in asthma: influences of genetics, environment, and T-cell plasticity. J Allergy Clin Immunol, 2013. 131(5): p. 1267–74; quiz 1275.

39. Newcomb, D.C. and R.S. Peebles, Jr., Th17-mediated inflammation in asthma. Curr Opin Immunol, 2013. 25(6): p. 755–60.

40. Finkelman, F.D., Anaphylaxis: lessons from mouse models. J Allergy Clin Immunol, 2007. 120(3): p. 506–15; quiz 516-7.

41. Speeckaert, R., et al., Th Pathways in Immune-Mediated Skin Disorders: A Guide for Strategic Treatment Decisions. Immune Netw, 2024. 24(5): p. e33.

42. Choy, D.F., et al., T_H_2 and T_H_17 inflammatory pathways are reciprocally regulated in asthma. Science Translational Medicine, 2015. 7(301): p. 301ra129–301ra129.

43. Brown, S.G., Cardiovascular aspects of anaphylaxis: implications for treatment and diagnosis. Curr Opin Allergy Clin Immunol, 2005. 5(4): p. 359–64.

44. Kounis, N.G., Kounis syndrome: an update on epidemiology, pathogenesis, diagnosis and therapeutic management. Clin Chem Lab Med, 2016. 54(10): p. 1545–59.

45. Simons, F.E., et al., World allergy organization guidelines for the assessment and management of anaphylaxis. World Allergy Organ J, 2011. 4(2): p. 13–37.

46. Sicherer, S.H. and H.A. Sampson, Food allergy: Epidemiology, pathogenesis, diagnosis, and treatment. J Allergy Clin Immunol, 2014. 133(2): p. 291–307; quiz 308.

47. Bischoff, S.C., et al., Intestinal permeability--a new target for disease prevention and therapy. BMC Gastroenterol, 2014. 14: p. 189.

48. Turner, J.R., Intestinal mucosal barrier function in health and disease. Nat Rev Immunol, 2009. 9(11): p. 799–809.

49. Liang, X., et al., Intravenously Administered Human Umbilical Cord-Derived Mesenchymal Stem Cell (HucMSC) Improves Cardiac Performance following Infarction via Immune Modulation. Stem Cells Int, 2023. 2023: p. 6256115.

50. Shao, L., et al., Inflammation in myocardial infarction: roles of mesenchymal stem cells and their secretome. Cell Death Discovery, 2022. 8(1): p. 452.

51. Jeong, S.Y., et al., Hyaluronic acid stimulation of stem cells for cardiac repair: a cell-free strategy for myocardial infarct. J Nanobiotechnology, 2024. 22(1): p. 149.

52. Bachiller, S., et al., Microglia in neurological diseases: a road map to brain-disease dependent-inflammatory response. Frontiers in cellular neuroscience, 2018. 12: p. 488.

53. Gieseck, R.L., M.S. Wilson, and T.A. Wynn, Type 2 immunity in tissue repair and fibrosis. Nature Reviews Immunology, 2018. 18(1): p. 62–76.

54. Buckley, C.D., et al., Endothelial cells, fibroblasts and vasculitis. Rheumatology (Oxford), 2005. 44(7): p. 860–3.

55. Papayannopoulos, V., Neutrophil extracellular traps in immunity and disease. Nature Reviews Immunology, 2018. 18(2): p. 134–147.

56. Berin, M.C. and H.A. Sampson, Food allergy: an enigmatic epidemic. Trends in immunology, 2013. 34(8): p. 390–397.

57. Voisin, T., A. Bouvier, and I.M. Chiu, Neuro-immune interactions in allergic diseases: novel targets for therapeutics. Int Immunol, 2017. 29(6): p. 247–261.

58. Modena, B.D., K. Dazy, and A.A. White, Emerging concepts: mast cell involvement in allergic diseases. Transl Res, 2016. 174: p. 98–121.

59. López-Fandiño, R., Role of human antigen-specific T cells in tolerance and IgE-mediated allergic reactions to food. Frontiers in Immunology, 2025. Volume 16 - 2025.

60. Alagappan, V.K., et al., Angiogenesis and vascular remodeling in chronic airway diseases. Cell Biochem Biophys, 2013. 67(2): p. 219–34.

61. Le, D.T., et al., Mismatch repair deficiency predicts response of solid tumors to PD-1 blockade. Science, 2017. 357(6349): p. 409–413.

62. Satija, R., et al., Spatial reconstruction of single-cell gene expression data. Nature biotechnology, 2015. 33(5): p. 495–502.

63. Stuart, T., et al., Comprehensive integration of single-cell data. cell, 2019. 177(7): p. 1888–1902. e21.

64. Hinton, G. and L. Van Der Maaten, Visualizing data using t-sne journal of machine learning research. Journal of Machine Learning Research, 2008. 9: p. 2579–2605.

65. Blondel, V.D., et al., Fast unfolding of communities in large networks. Journal of statistical mechanics: theory and experiment, 2008. 2008(10): p. P10008.

66. Aran, D., et al., Reference-based analysis of lung single-cell sequencing reveals a transitional profibrotic macrophage. Nature immunology, 2019. 20(2): p. 163–172.

67. Yang, A.C., et al., Dysregulation of brain and choroid plexus cell types in severe COVID-19. Nature, 2021. 595(7868): p. 565–571.

68. Guo, J., et al., Single-cell transcriptomics in ovarian cancer identify a metastasis-associated cell cluster overexpressed RAB13. J Transl Med, 2023. 21(1): p. 254.

