## Supplementary figure 2 and supplementary figure 6 for "A Multi-Organ Single-Cell Atlas Maps Allergen and Organ-Specific Cellular Dynamics in the Progression from Food Allergy to Anaphylactic Shock"

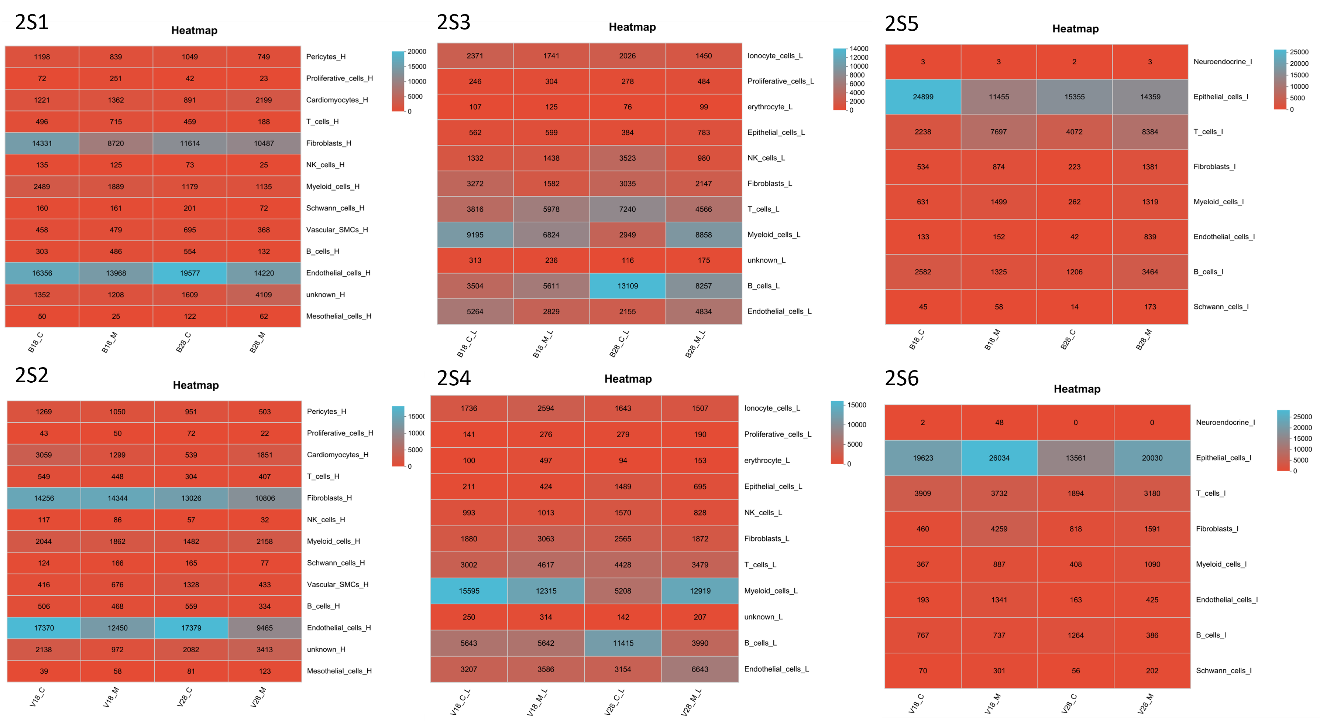

Fig 2S: Heatmap representation of change in cell number in experimental animals. Heatmap color represents cell number in heart (2S1,2S2), lung (2S3,2S4) and intestine (2S5,2S6) in OVA and BSA model. X-axis in the heatmap represents BSA (B) or OVA (O), phase I (18) or phase II (28) and control (c) or model (M). Y axis represents the cell type in heart (H), lung (L) and intestine (I).

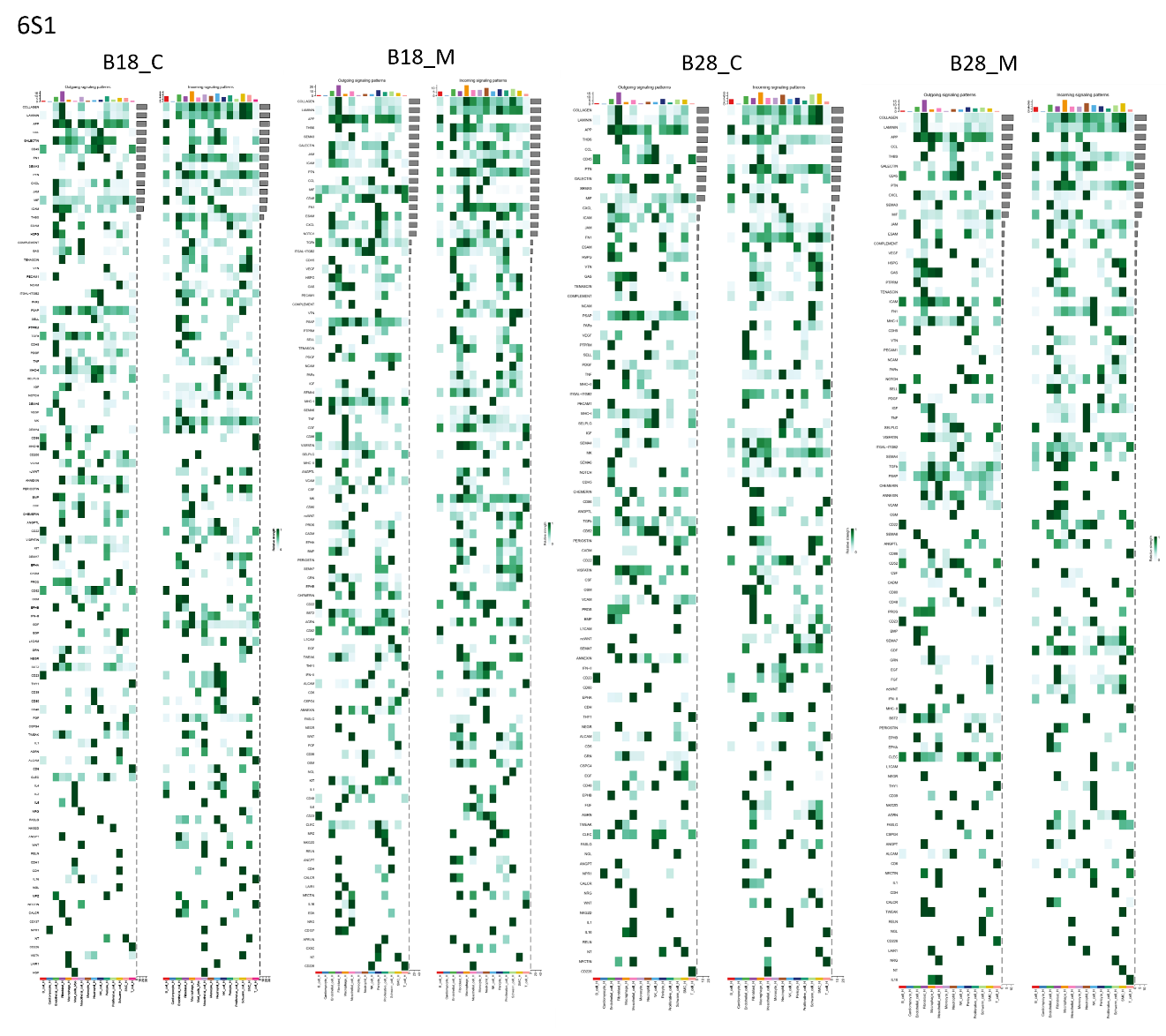

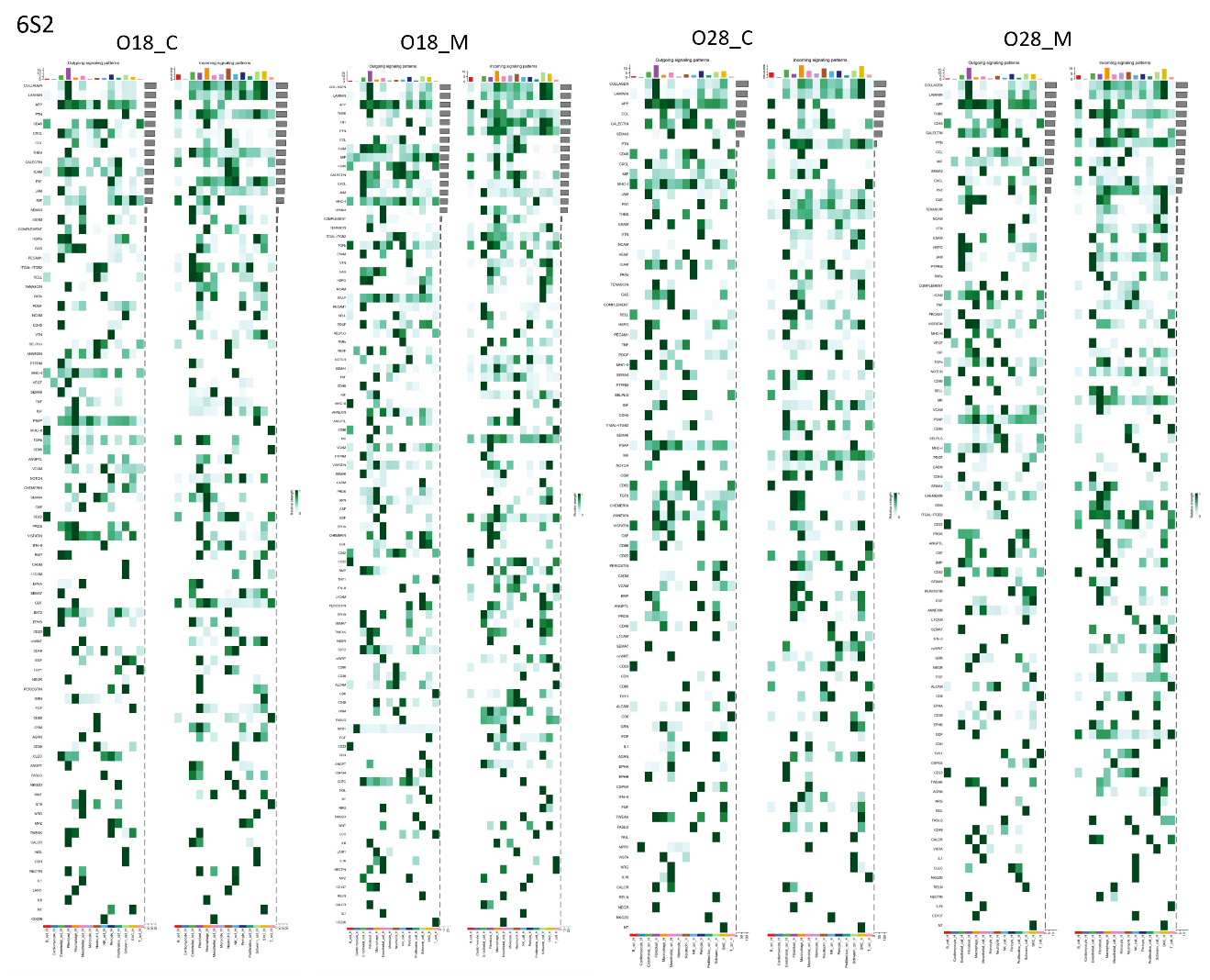

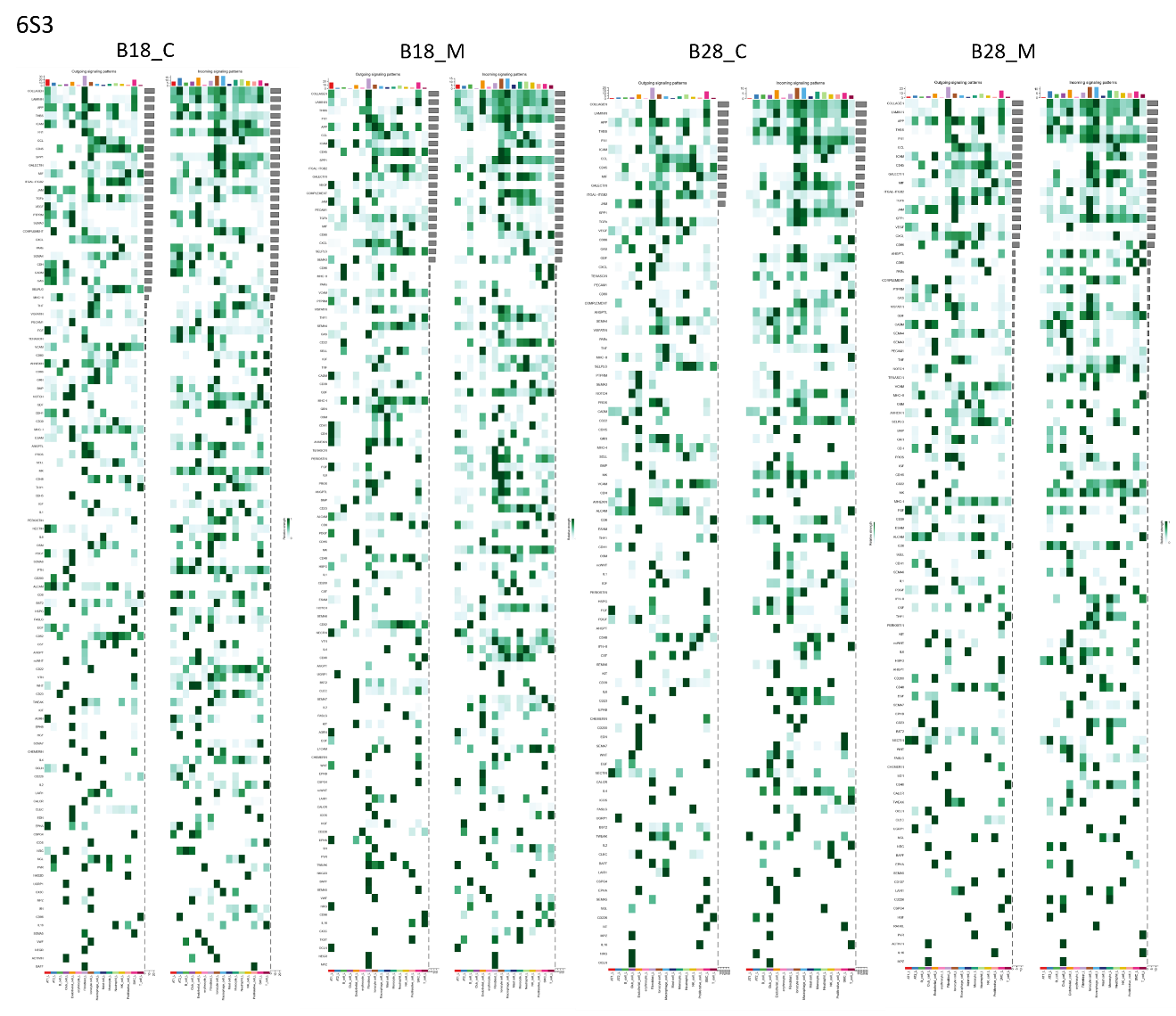

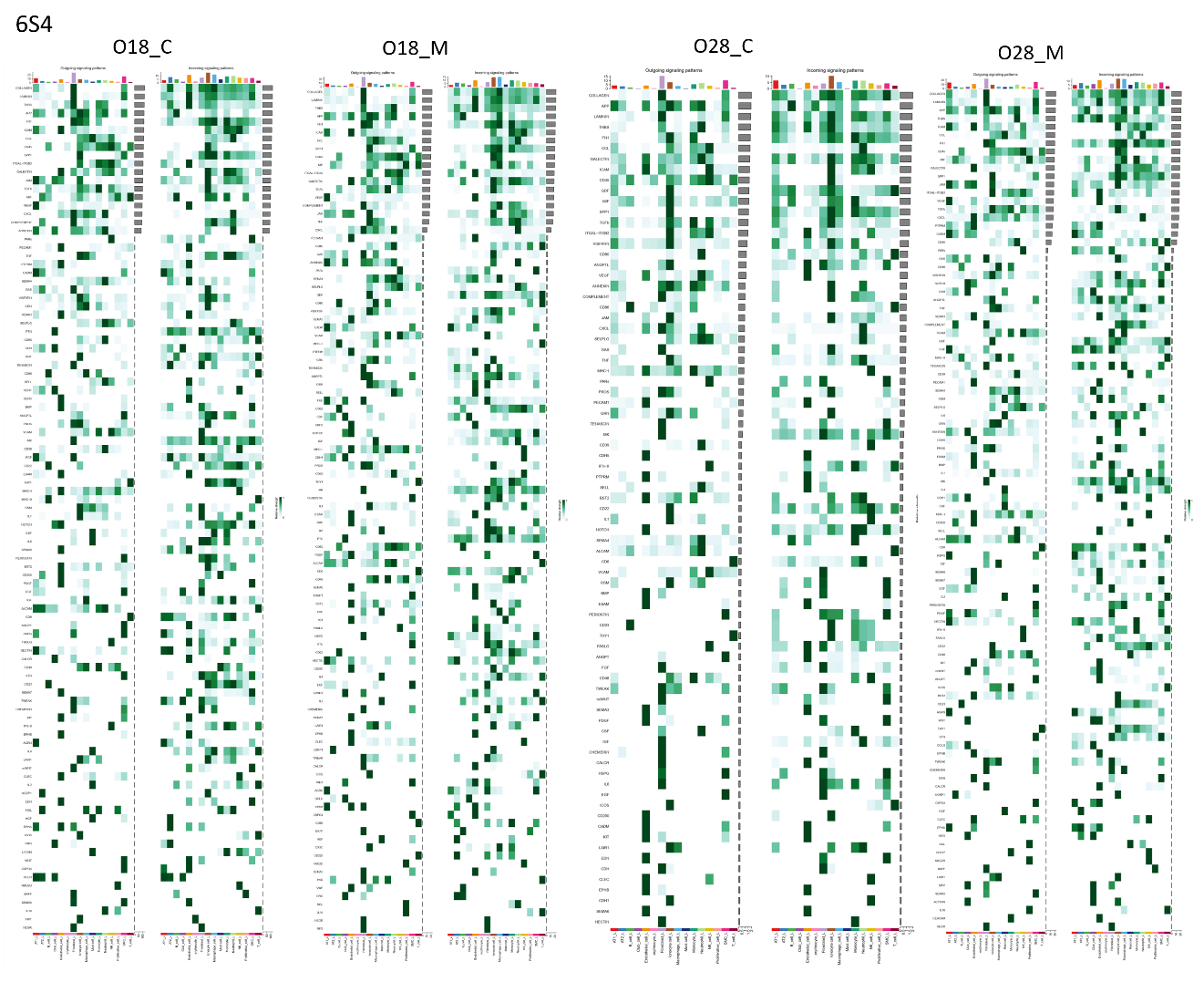

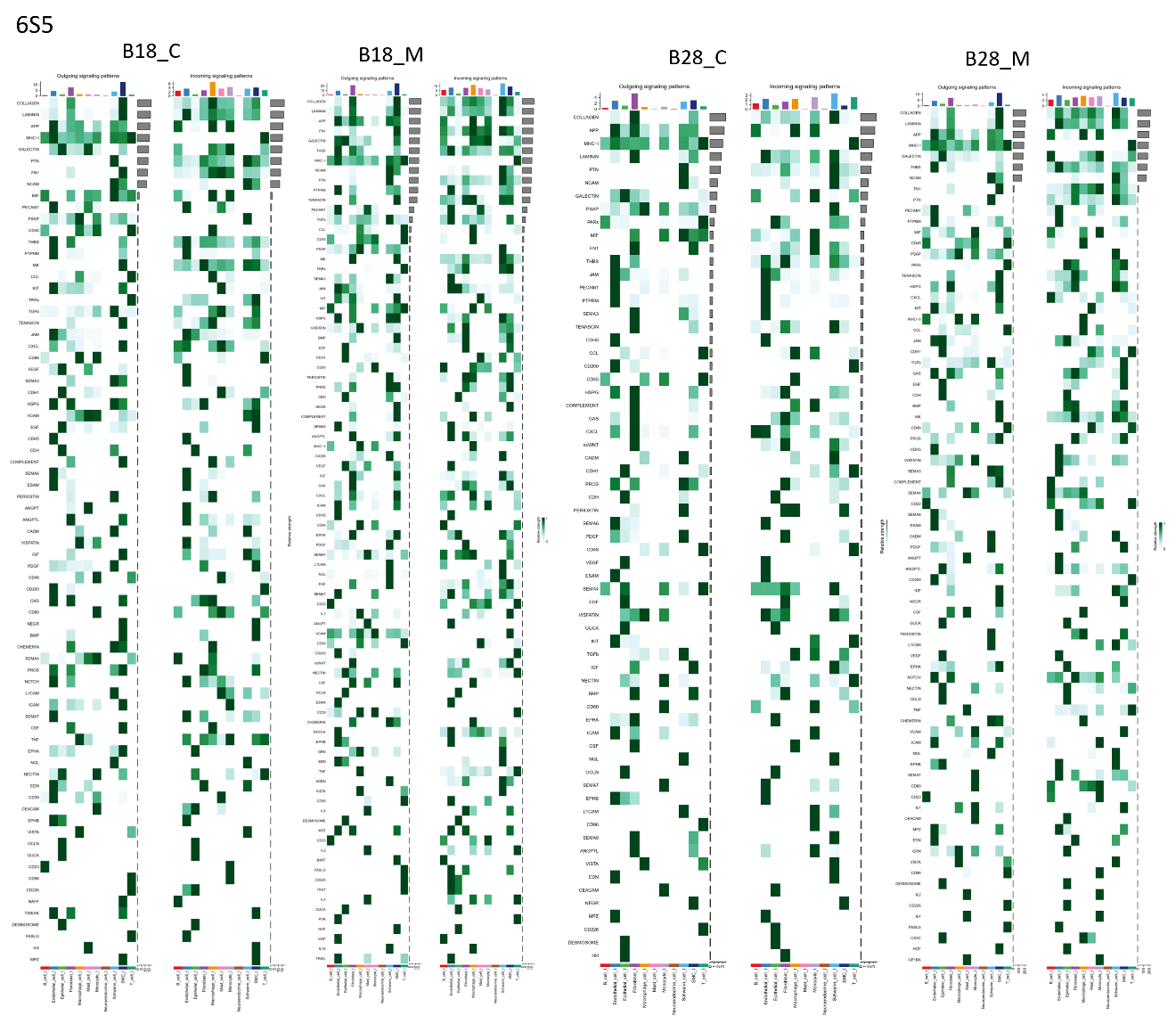

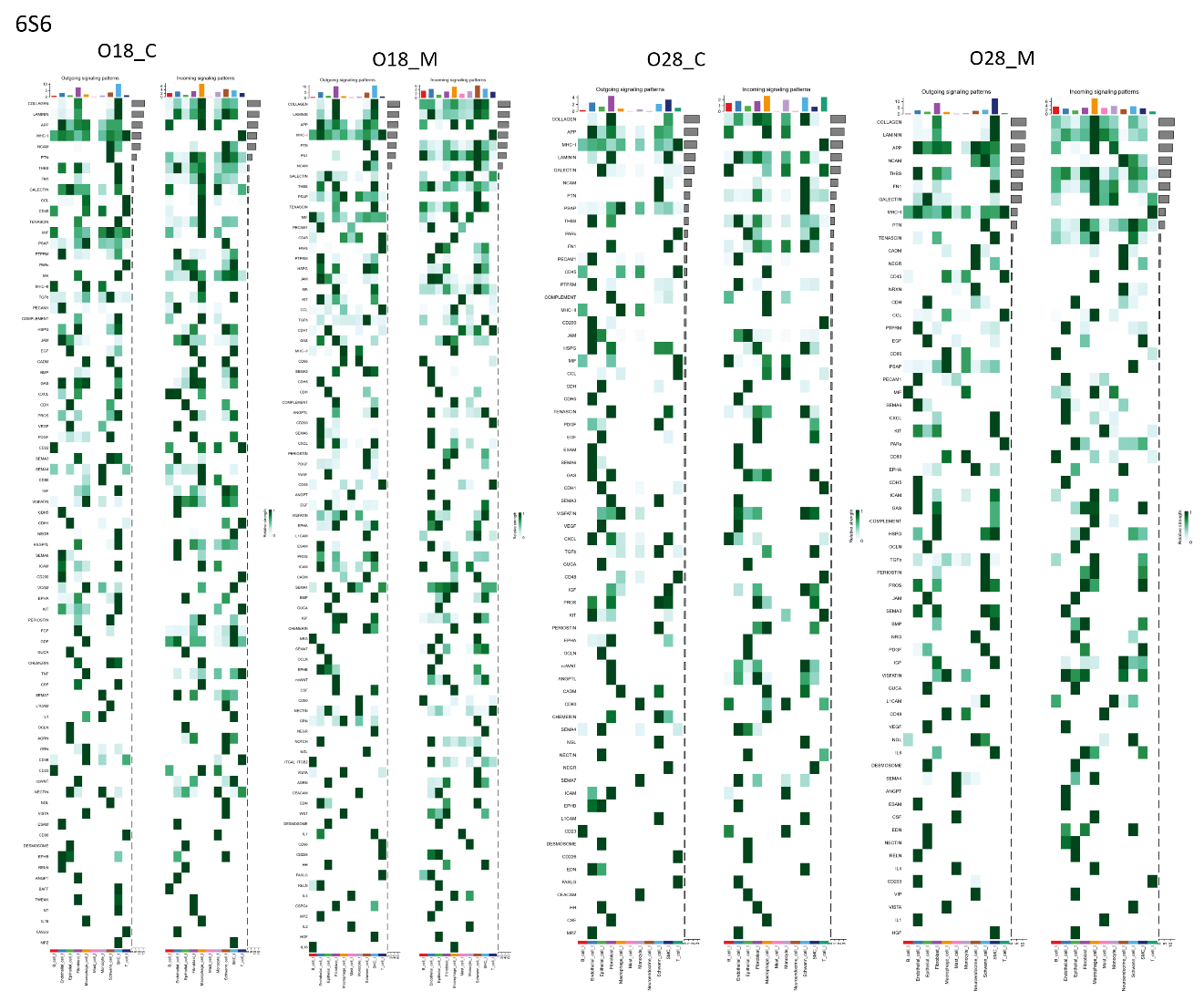

**Fig 6S: Heat map representation of the signal flow pattern between the cells using CellChat communication analysis.** Each figure is classified into incoming and outgoing categories for BSA and OVA models for phase I (day 18) and phase II (day 28). Incoming and outcoming signal flow pattern in the cells for heart (6S1,6S2), lung (6S3, 6S4) and intestine (6S5,6S6). The heatmap color bars show the corresponding signal strength of signaling pathways in different cell groups. The colored bar graph at the top shows the total signal intensity of a cell population by summarizing all of the signaling pathways shown in the heat map, while the gray bar graph on the right shows the overall signal magnitude for a signaling pathway by summarizing all of the cell populations shown in the heat map. Facet label above heatmap represents BSA (B) or OVA (O), phase I (18) or phase II (28) and control (c) or model (M).
